# Integrated *in silico* and experimental screening identifies novel ligands that target precursor microRNA-31 at the Dicer cleavage site

**DOI:** 10.1101/2025.06.20.660744

**Authors:** Grace Arhin, Lily Haghpassand, Sarah C. Keane

## Abstract

MicroRNAs (miRNAs) regulate gene expression and the dysregulation in mature miRNA levels has been implicated in a wide variety of diseases. In particular, altered levels of mature microRNA-31 (miR-31) has been linked to a variety of different cancers. Targeting functionally relevant sites of the precursor structure of miR-31 with small molecules offer a strategy to regulate miR-31 maturation. Herein we describe a virtual screening approach to explore the druggability of the precursor structure of microRNA-31 (pre-miR-31). We used a structure-guided approach to virtually screen a fragment library and followed up with experimental characterization of top-ranking candidates, leading to the identification of several compounds that bound to pre-miR-31. Further characterization of the RNA-ligand complexes by heteronuclear single quantum coherence (HSQC) NMR spectroscopy revealed three compounds bound pre-miR-31 at the Dicer cleavage site, suggesting that these compounds may function to inhibit Dicer processing. Using these initial hits, we performed chemical structure similarity searches and identified additional binders of pre-miR-31 that had equivalent or enhanced binding relative to the parent compounds. These studies suggest a generalizable approach by which RNA-binding ligands can be identified from large chemical databases. These hits can then be further optimized to improve affinity and specificity for downstream functional assays.

## Introduction

Over the past two decades there has been an exponential growth in our understanding of non-coding (nc) RNAs and their functions in a wide range of human diseases.^1–3^ These discoveries have paved the way for ncRNAs to be recognized as viable therapeutic targets.^4,5^ Among the extensively studied ncRNAs are microRNAs (miRNAs) and they play crucial roles in disease development and progression.^3,6,7^ Mature miRNAs function by targeting complementary sequences within messenger RNAs (mRNAs) for translational repression.^8,9^ Initially transcribed as long hairpin structures (primary (pri-) miRNAs),^10^ miRNAs mature through two distinct enzymatic cleavage steps. Pri-miRNAs are cleaved by the Drosha in complex with DiGeorge syndrome critical region 8 (DGCR8) into a shorter hairpin precursor (pre-) miRNAs.^11–14^ Dicer/TRBP then cleaves the pre-miRNA to generate mature miRNAs.^15–17^ Dysregulation in the mature levels of miRNAs causes disease progression, in part due the role that miRNAs play in post-transcriptional regulation.^7,18,19^

The structure of precursor microRNA-31 (pre-miR-31) was recently determined by nuclear magnetic resonance (NMR) spectroscopy.^20^ This structure revealed the presence of a triplet of base pairs (junction) that connected its apical loop to a bulge at its Dicer/TRBP cleavage site (dicing site). Through mutagenesis studies, the stability of the junction base pairs was found to impact Dicer/TRBP cleavage, affecting the levels of mature miR-31 produced *in vitro.*^20^ Interestingly anti sense oligonucleotides (ASOs) designed to target the pre-miR-31 structure was found to inhibit Dicer/TRBP cleavage in vitro and in cells.^21^ Dysregulation in the levels of mature miR-31 has been linked to multiple different cancers including colorectal cancers and cervical cancers, highlighting the importance of regulating their levels in cells.^22,23^ Although not a validated drug target, a promising strategy to modulate mature miR-31 levels and potentially its function in diseases could involve using small molecules to alter the pre-miR-31 structure and dynamics which has been shown to influence its maturation by Dicer/TRBP.^20^

Herein we describe the implementation of an integrative approach to rapidly explore the ligandability of the pre-miR-31 structure. Such an assessment would not only provide foundational insights into pre-miR-31-small molecule interactions but also guide the rational design of more potent and perhaps more selective compounds for downstream applications. To streamline this effort, we employed a multi-assay screening strategy which began with a computational library screen, followed by two independent binding validation assays to optimize hit detection. Confirmed hits were then used for chemical structure similarity searches on commercial databases to identify additional small molecule binders of the miR-31 hairpin.

Using this optimized integrative screening approach, we identified several unique small molecules that bound pre-miR-31. Our model of the RNA-small molecule complexes, informed by heteronuclear quantum coherence (HSQC-) NMR, revealed that these ligands bind at the dicing site of the pre-miR-31 potentially inducing a destabilization that disrupts the base pairs at the junction region.

## Results and Discussion

### In silico screening of NCI Diversity set IV library to identify small molecule binders of pre-miR-31

The 3D ensemble of pre-miR-31 revealed a well-defined apical loop and junction base pairs connecting the bulge at the dicing site to the apical loop (**Fig. 1a, b**).^20^ We performed virtual screening to rapidly identify potential binders of the hairpin. Briefly, we first identified ligandable cavities within the pre-miR-31 NMR ensemble using RNACavityMiner.^4^ Through this analysis, we observed that across the entire NMR ensemble, the major groove near the junction residues exhibited highest ligandability score. Additionally, we found that for some conformers (conformers, 4, 10, 12, 15), RNACavityMiner predicted higher ligandability scores for major grooves around the C•A and G•A mismatches (**Fig. S1**). Next, a total of 1,596 small molecules from the National Cancer Institute’s (NCI) developmental therapeutics (DTP) library were screened against the ligandable pockets identified within the 3D NMR ensemble of pre-miR-31 using our previously optimized virtual screening protocol^24^ (**Fig. 1c**) with a few modifications. Details of this screening protocol are highlighted in **Fig. S2**. Initial hits (53) were identified and of these, we acquired 40 compounds with the lowest (most energetically favored) docking score from the NCI DPT for binding characterization. The chemical structures of all 40 compounds are shown in **Figs. S3 - S5** highlighting their NCI compound identity number as well as their assigned names which has been used for ease and clarity throughout the results and discussion.

**Figure 1:**
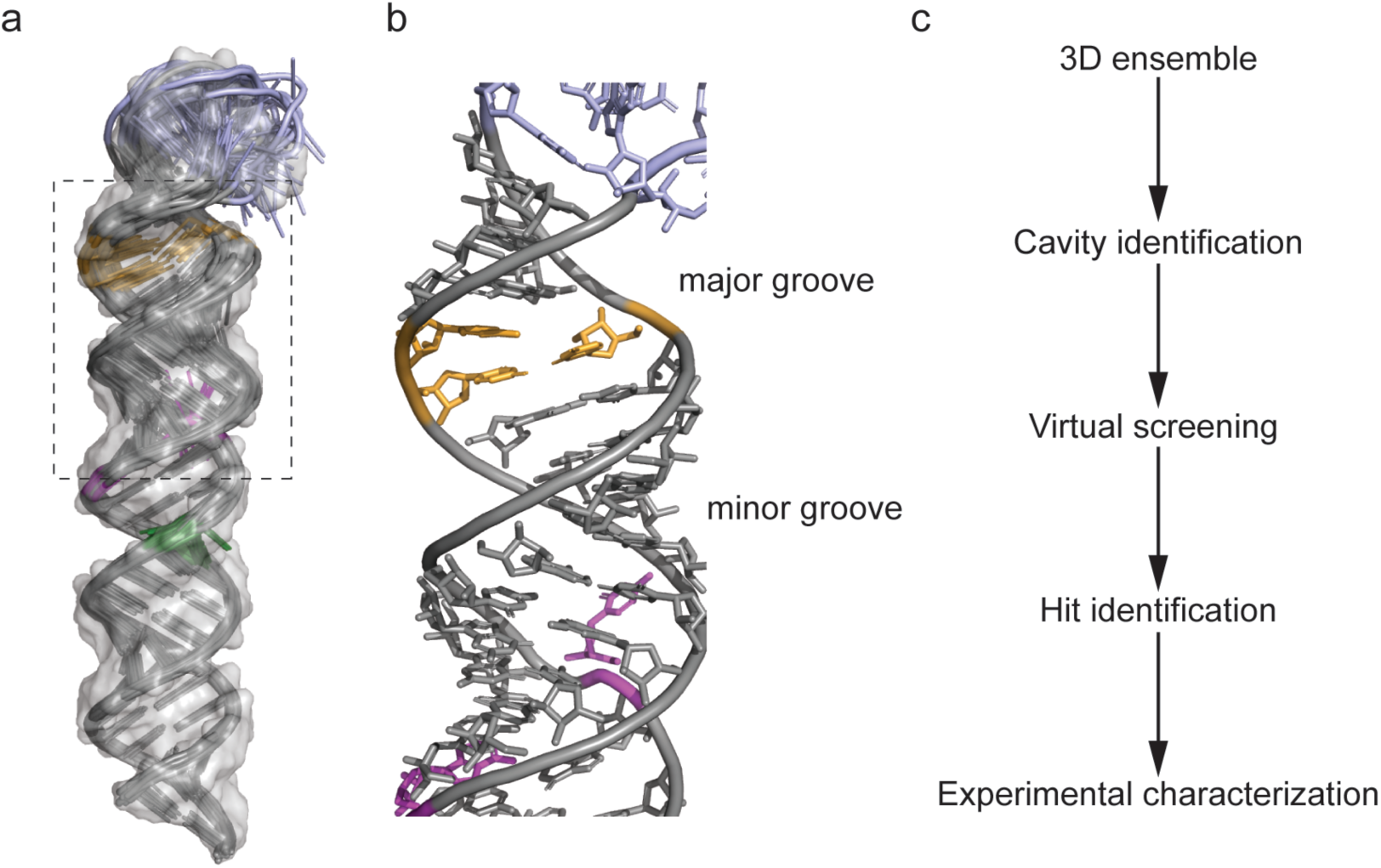
NMR ensemble based virtual screening. (a) NMR ensemble of pre-miR-31 highlighting the apical loop residues in blue, dicing site residues in orange, C•A mismatch in purple and G•A mismatch in green with a transparent surface rendering. (b) Enlarged view of residues highlighted with a black box in panel a. (c) Ensemble based virtual screening workflow.

### Dual binding detection by STD-NMR and fluorescent indicator displacement assay

Based on our previous studies, which identified important structural features within the pre-miR-31 that influence Dicer/TRBP cleavage,^20^ we reasoned that the most significant effects would be small molecules that target the dicing site or dynamics at the junction region of the RNA. Notably our cavity searching algorithm on the entire ensemble of pre-miR-31 shows a high ligandability score for this region (**Fig. S1**) across the ensemble. We therefore chose to experimentally assay ligands using a pre-miR-31 construct that spans residues 20 through 52 (**Fig. 2a**) which for the rest of the discussion is referred to as the miR-31 hairpin. We validated our *in-silico* hits experimentally using two independent screening assays: saturation transfer difference (STD) NMR spectroscopy and a fluorescence indicator displacement (FID) assay. This dual screening approach is valuable because it combines the unique strengths of two distinct techniques.

**Figure 2:**
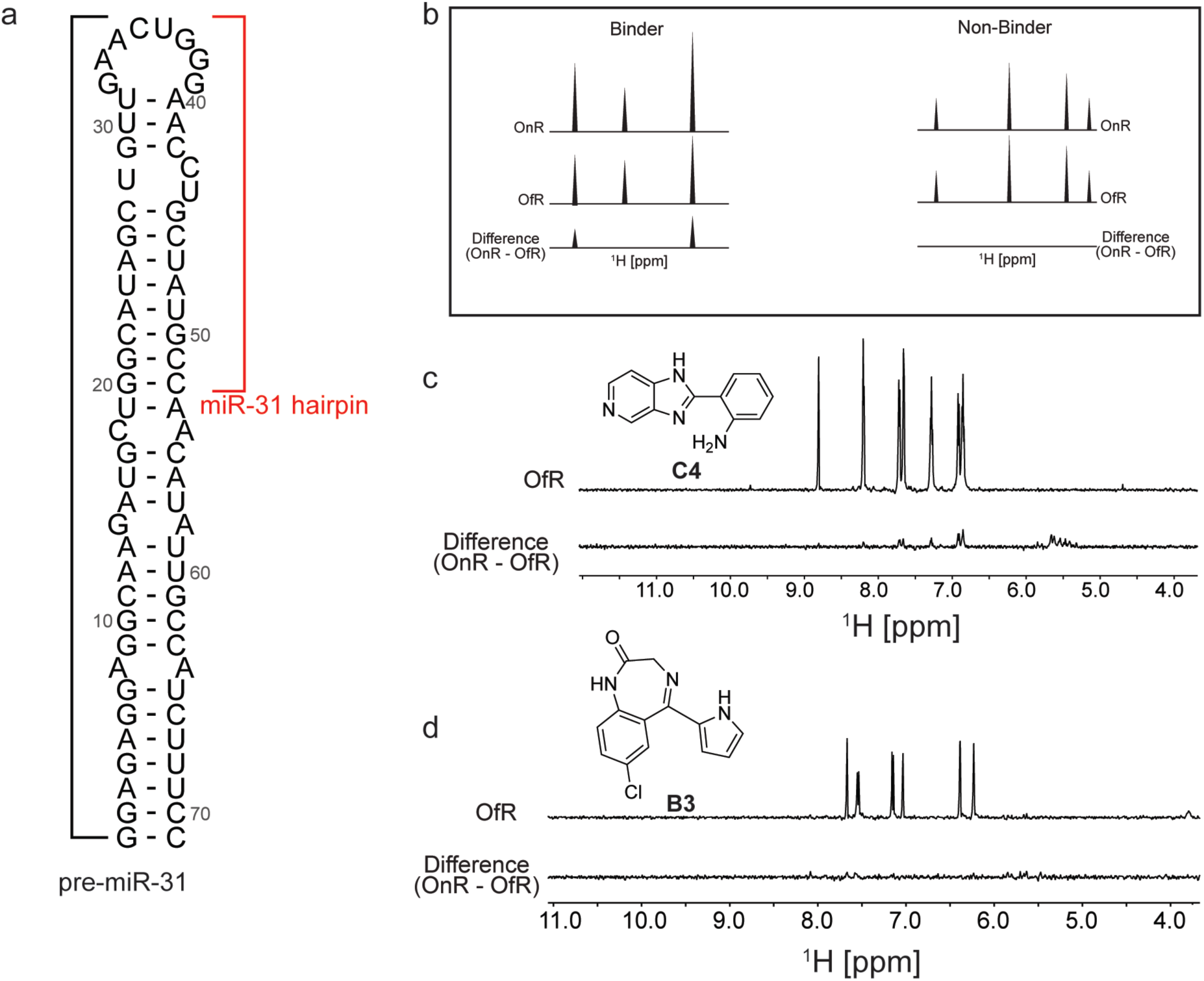
Identification of small molecule binders of the miR-31 hairpin by STD-NMR. (a) Secondary structure of pre-miR-31 highlighting the miR-31 hairpin used in study. (b) Criteria for binding selection in STD-NMR experiments. (c-d) Representative spectra of (c) a miR-31 hairpin binder and (d) a miR-31 hairpin non-binder.

STD-NMR spectroscopy allows monitoring the changes in a small molecule proton spectrum when bound by a biomolecule.^25^ This method has been previously employed to characterize several RNA-small molecule interactions.^24,26,27^ We began by acquiring the ^1^H-NMR spectrum of our identified *in silico* hits at ∼500 μM in NMR buffer [50 mM KH2PO4 at pH 7.5, 1 mM MgCl2] containing 2% DMSO and 10% D2O. We observed that roughly half (22/40) of the compounds had detectable proton signals and as such were suitable for binding characterization by STD-NMR spectroscopy (**Figs. S6 – S27**). Next, we acquired the STD-NMR spectra for all 22 compounds at a 1:20 RNA:ligand ratio (10 μM RNA:200 μM ligand). The criterion for hit selection is illustrated in **Fig. 2b**. A ligand was considered a binder if proton signals were observed in its difference spectrum and a non-binder if no proton signals were detected in its difference spectrum. After analyzing the difference spectrum of all 22 compounds, we identified 7 compounds (A1, A5, A9, B7, C4, C6, and RA3) that bound the miR-31 hairpin (**Fig. 2c and Fig. S28**). The remaining 15 compounds had no detectable proton signals in their difference spectra and were considered non-binders. A representative spectrum of one non-binder from our testing library is shown (**Fig. 2d**).

Simultaneously, we tested the ability of the *in silico* ‘hits’ to displace TO-PRO-1 dye in an RNA-dye complex (**Fig. 3a**). This approach offers one advantage relative to STD-NMR spectroscopy because, in theory, there is no limitation to the type of ligand that can be analyzed. FID assays have been successfully employed to screen for RNA binding small molecules.^28,29^ We fine-tuned the assay to our RNA by determining the affinity of TO-PRO-1 for the miR-31 hairpin to determine the concentration of the miR-31 hairpin bound by TO-PRO-1 at an initial fraction of 0.1 (fb0.1) which is important to improve assay sensitivity (**Fig. S29a**).^30,31^ Our small molecules were screened at 100 μM concentration and hits were defined as small molecules that displaced TO-PRO-1 by more than 15% in duplicate experiments. Out of the 40 compounds tested, we observed significant reduction in fluorescence in the RNA-dye complex upon incubation with four compounds (DA1, A1, C11 and DA2), indicative of binding (**Fig. 3b**). 25 compounds showed no significant change in the RNA-dye fluorescence (**Fig. 3b**). For the remaining 11 compounds (A10, A11, B1, B5, B11, C1, C3, C6, C7, RA1 and RA4) we observed significant increase in the RNA-dye fluorescence upon their addition (**Fig. S29b**).

**Figure 3:**
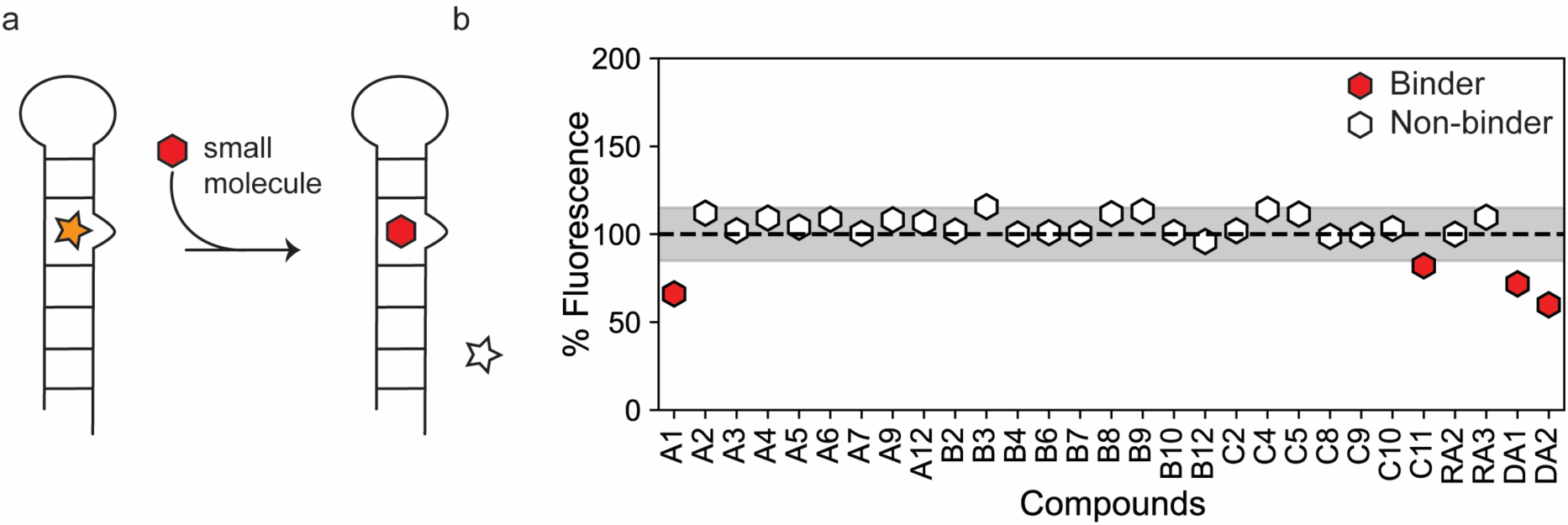
Identification of small molecule binders of the miR-31 hairpin using an FID assay. (a) Schematic of the FID assay using TO-PRO-1. TO-PRO-1 fluoresces upon binding the RNA (orange star). When a competitive small molecule (red hexagon) displaces bound TO-PRO-1, fluorescence is decreased (clear star) (b) Percentage fluorescence change in RNA-dye complex by small molecules. The figure includes a line at 100% representing an ideal scenario where there is no change in fluorescence. The shaded region of ± 15% denotes insignificant fluorescent changes to avoid false positives. Ligands showing a fluorescence change >15% (<85%) were considered hits.

To further interrogate potential interactions between the ligands, TO-PRO-1, and/or the RNA, we measured the fluorescence when TO-PRO-1 was incubated with the small molecule ligands or when the RNA was incubated with the small molecule ligands. We observed that the addition of compounds A9, A10, A11, B6, B9, B12, C1, C3, C6, and C11 to TO-PRO-1 led to fluorescence increase when compared to TO-PRO-1 in buffer alone (**Fig. S29c**). The findings suggest that these ligands interact with TO-PRO-1 which accounts for the fluorescence increase observed for A10, A11, C1, C3, and C6 when added to the RNA-dye complex. Similarly, addition of A10, B12, C1, and C3 to the RNA led to an increase in fluorescence when compared to RNA in buffer alone (**Fig. S29c**).

For comparison, we show a summary of the results for the two screening approaches (**Fig. S30**). We observed binding for A1, A5, A9, B7, C4, C6, and RA3 compounds using our STD-NMR screening approach and A1, C11, DA1, and DA2 compounds using the fluorescence indicator displacement assay. We had no STD-NMR data for C11, DA1, and DA2 because these compounds had no detectable ^1^H spectra and therefore cannot assess binding by STD-NMR. However, it is evident that our FID assay could not detect binding interactions for A5, A9, B7, C4, C6, and RA3, counter to what we expected based on the STD-NMR analysis. Ligand observed NMR experiments are well known for their ability to detect very weak interactions between a biomolecule and a small molecule.^32^ On the contrary, an indicator displacement assay might require a compound that binds with moderate affinity to be able to displace the dye due to the strong affinity between the dye and the RNA. These findings demonstrate the importance of combining different approaches for ligand screening because a single assay may result in loss of information. Collectively, we validated binding of 25% of the compounds identified in our *in silico* screen.

### C4, A1 and DA1 induce perturbations in the miR-31 hairpin at the junction region and dicing site

To precisely locate the small molecule ligand interaction site on the miR-31 hairpin, we acquired a ^1^H-^13^C HSQC spectrum of ^13^C/^15^N-A/C-labelled miR-31 hairpin in the presence of ligands identified by our experimental screen. Our isotope labelling approach enabled high sequence coverage while minimizing signal overlap in the spectrum. Because the ligands were dissolved in DMSO, we first acquired the HSQC spectrum of the miR-31 hairpin at 0% and 2% DMSO. These spectra revealed very subtle changes in the RNA resonances, indicating that 2% DMSO did not significantly disrupt the RNA structure (**Fig. S31**). For single-point titrations, ligands were present in 10-fold excess relative to RNA concentration.

Interestingly, compounds C4, A1, and DA1 induced very significant perturbations in the miR-31 hairpin. The largest perturbations were localized in the residues at the junction region and dicing site of the miR-31 hairpin for C4 (**Fig. 4a-c)** and A1 (**Fig. S32**). These perturbations included the C2-H2 correlations of residues A40 and A41 as well as C6-H6 correlations of residues C42 and C43. For DA1, perturbations were observed across all resonances of the miR-31 hairpin in the presence of 10-fold excess of the ligand (**Fig. S33**). In a 1:1 titration however, these perturbations were localized to the C2-H2 correlations of A40, A41 and C6-H6 correlations of C42 and C43 (**Fig. S34**), reflecting the non-specific nature of ligand binding in the presence of excess DA1. This finding additionally suggests that DA1 binds the miR-31 hairpin with a higher affinity, compared to A1 and C4.

**Figure 4:**
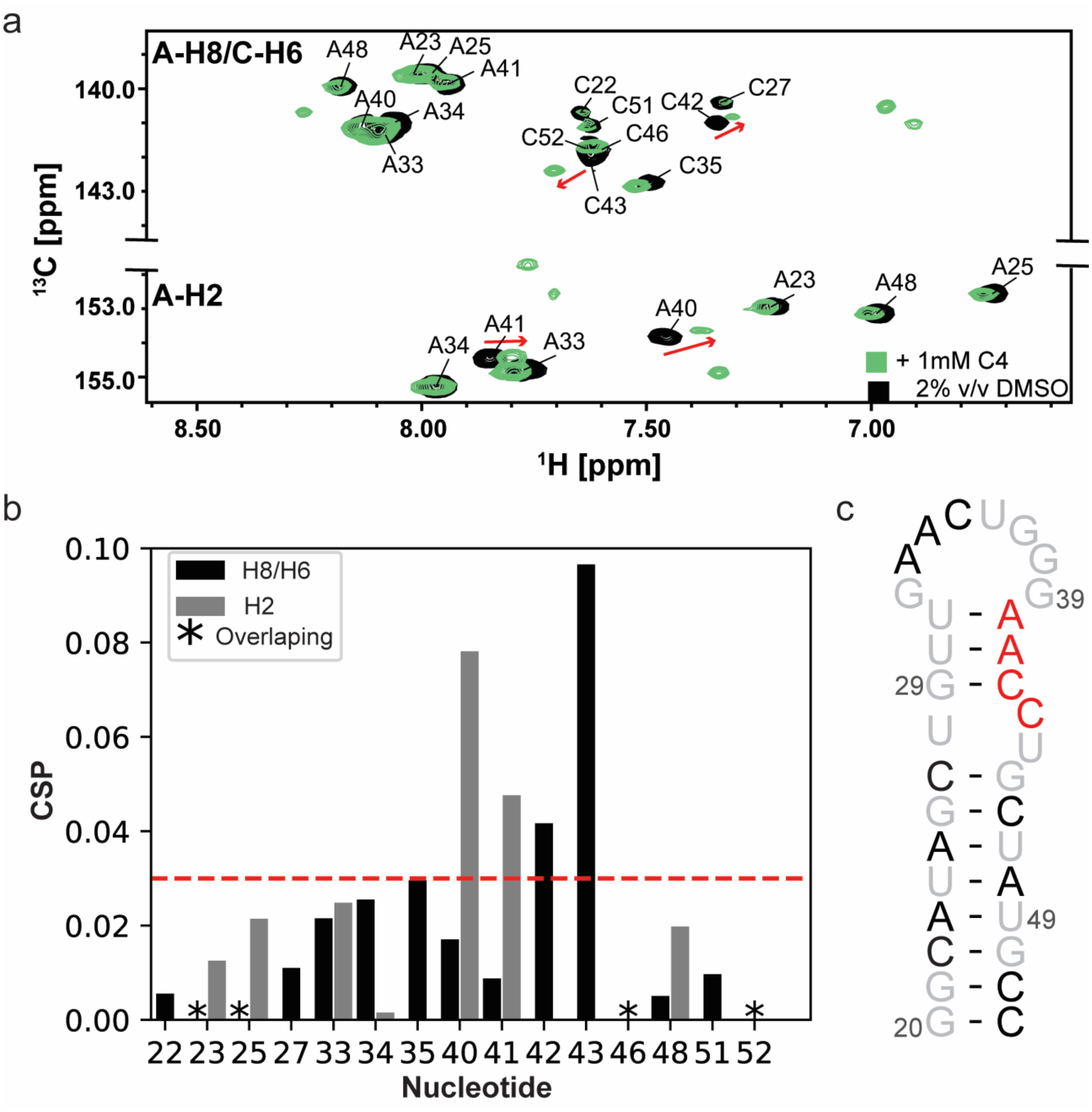
C4 induced changes in the miR-31 hairpin. (a) Overlayed ^1^H-^13^C HSQC spectra of ^13^C/^15^N adenosine cytosine labelled miR-31 hairpin in 2% (v/v) DMSO (black) and C4 (green). (b) Calculated chemical shift perturbation (CSPs) values induced by addition of C4 to the miR-31 hairpin. Gray bars indicate the perturbations in the C2-H2 correlations, black bars indicate perturbations in C6-H6/C8-H8 correlations. Black asterisks indicate overlapping signals that could not be quantified. (c) The most significant perturbations are mapped to the junction region of the miR-31 hairpin (red). Black residues were not significantly perturbed and grey residues were not

Surprisingly, the remaining compounds induced very subtle to no perturbations in the miR-31 hairpin spectrum (**Figs. S35 – S38**). The absence of chemical shift perturbations for the RNA in the presence of 10-fold excess of ligand is indicative of a very weak interaction.

### Chemical structure similarity search enables the discovery of additional miR-31 binders

Similarity searching is founded on the principle that if two small molecules have similar structures, they are likely to exhibit comparable properties. In our instance the property being examined was their binding interactions with the miR-31 hairpin. Having identified 3 unique scaffolds that interact at a functionally relevant site of pre-miR-31 (**Fig. 5a-c**), our objective was to investigate whether derivatives of our initial scaffolds would also bind the miR-31 hairpin. This similarity search relies on 2D structures rather than 3D conformations of the small molecules, thus the precise 3D pose of the ligand is not required for such analysis.^33^ This approach is especially useful when there is no experimentally determined ligand pose to be used in a pharmacophore-based screening. We identified five commercially available derivatives of our initial scaffolds (**Fig. 5d-g**). First, we docked the derivative compounds into the identified binding pocket of pre-miR-31 and compared their binding energies to those of the original scaffolds. As expected, most derivatives had similar docking score to their parent compounds (**Fig. 5a-f**), except for C4A which had a much lower (i.e. more energetically favorable) docking score compared to C4 (**Fig. 5c**,**g**).

**Figure 5:**
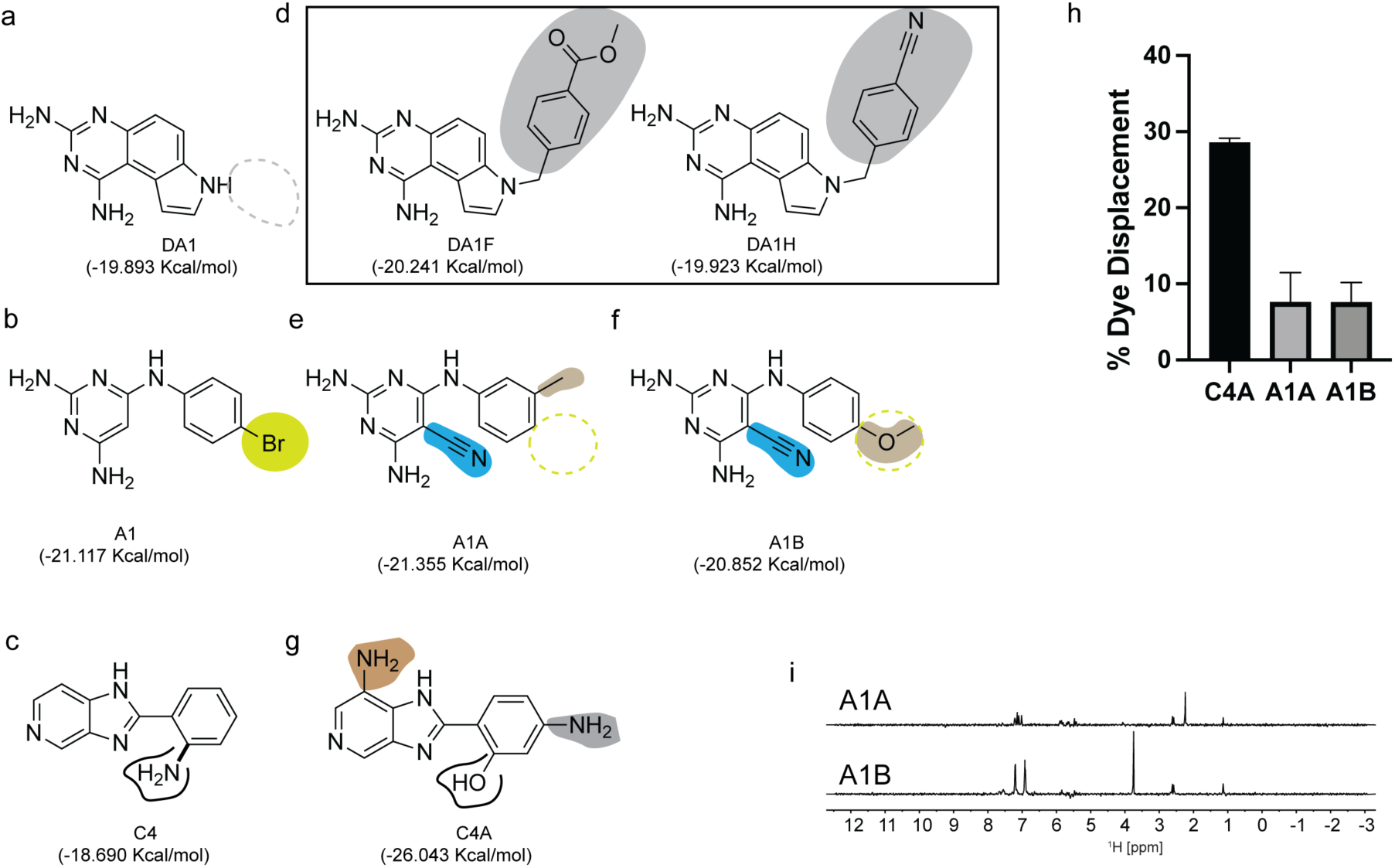
Chemical structure similarity search for additional binders of the miR-31 hairpin. (a-c) Chemical structures of (a) DA1, (b) A1, (c) C4 with their predicted docking scores. (d-g) Commercially available derivatives of (d) DA1, (e-f) A1 (g) C4 highlighting their differences relative to their parent scaffold along with their computed docking scores. (h) Plot of %TO-PRO-1 displaced for evaluation of binding of new derivatives to the miR-31 hairpin. (i) STD-difference spectra of derivatives A1A (top) and A1B (bottom).

Next, we sought to determine binding to the miR-31 hairpin *in vitro*. Although compound DA1 may bind the pre-miR-31 hairpin more tightly than A1 and C4, based on our HSQC experiments, we chose to focus on A1 and C4 derivatives instead. DA1 exhibited poor solubility in our aqueous buffer and inspection of the derivative structures, were likely to be even less soluble. Thus, we purchased A1A, A1B and C4A for binding characterization. First, we tested the ability of compounds A1A, A1B and C4A to displace TO-PRO-1 using our FID assay. Notably C4A displaces TO-PRO-1 at about 28%, indicating a binding interaction with the miR-31 hairpin. However, A1A and A1B displaced TO-PRO-1 at about 9% (**Fig. 5h**) which didn’t meet our previously described criteria for binding (**Fig. 3**). Nonetheless we observed binding by STD-NMR characterization (**Fig. 5i**), further supporting the notion that NMR can detect very weak binding events. We can therefore argue that in our second screening approach by similarity searching we had a 100% success at hit identification although we had a very small sample size.

### C4A binds the miR-31 hairpin with > 20-fold affinity than C4

To assess whether the hit derivatives had altered binding affinities relative to the parent compounds, we performed titration experiments to measure the affinities. We acquired a series of HSQC experiments where the ligand concentrations were increased while the RNA concentration remained constant and monitored perturbations in the RNA chemical shifts. We observed that addition of 10-fold excess of A1A and A1B induced insignificant perturbations in the RNA, suggesting weaker interaction with the miR-31 hairpin relative to the parent compound, A1 (**Fig. S39**).

Interestingly, in our analysis of C4 and C4A, we observed that addition of as low as 10 μM C4A (1:10 ligand:RNA molar ratio) induced line broadening of the A40 C2-H2 and C42 C6-H6 resonances with almost complete peak disappearance at around 50 μM C4A (indicative of saturation) with an estimated KD of 29.10 ± 0.15 μM (**Fig. 6a-b**). We note however that once the binding is at saturation at the binding site, addition of excess ligand led to non-specific binding interactions across the entire RNA. In contrast C4 did not reach saturation even at 1 mM concentration (**Fig. 6c-d**). These results suggest that the C4 scaffold could be derivatized, and that one such compound (C4A), bound the miR-31 hairpin with higher affinity.

**Figure 6:**
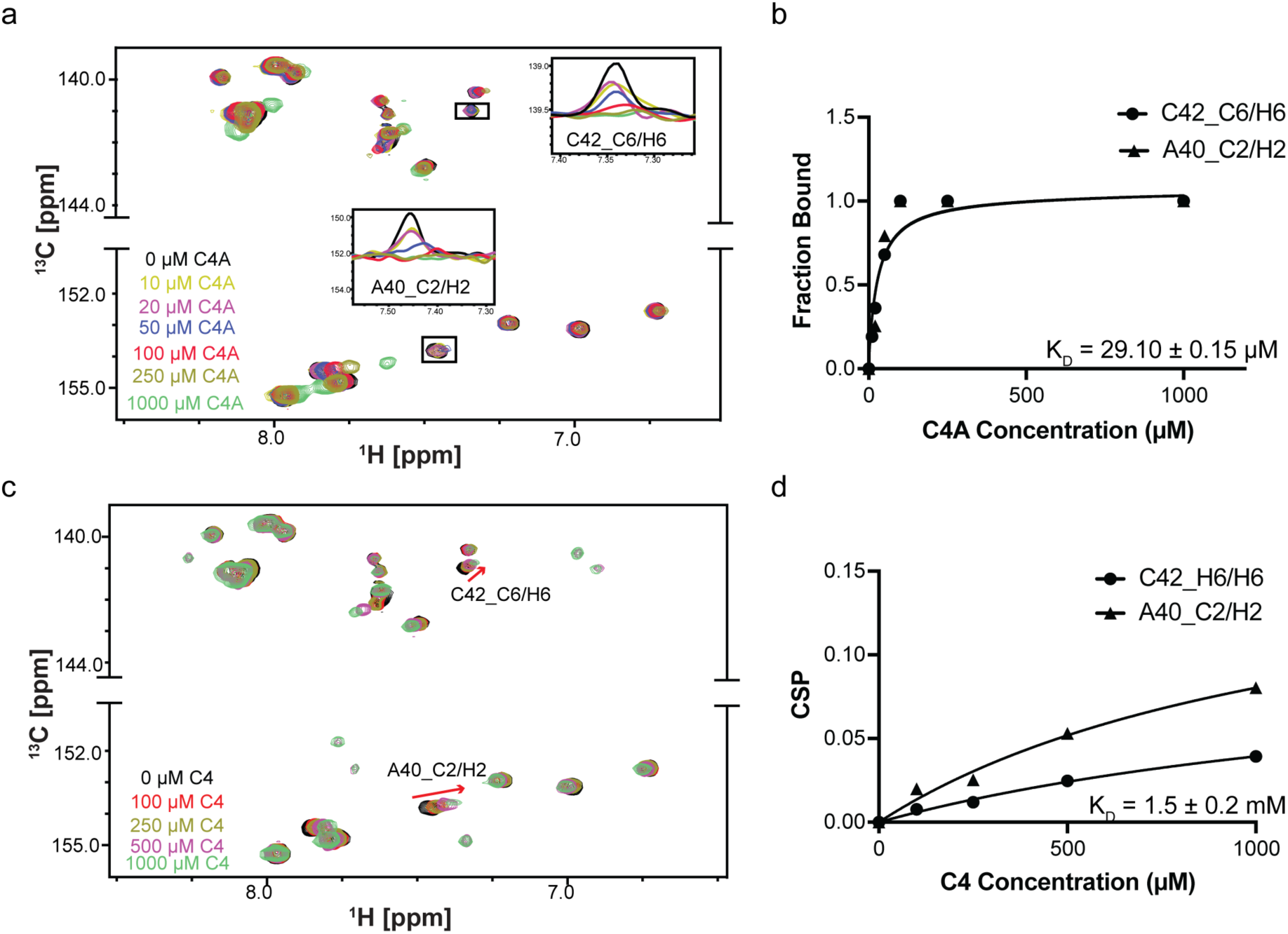
Binding affinities of C4A and C4. (a) Overlayed ^1^H-^13^C spectra of miR-31 hairpin at different C4A concentrations. (b) Fits of the chemical shift perturbations as function of C4A concentration (panel a). (c) Overlayed ^1^H-^13^C spectra of miR-31 hairpin at different C4 concentrations. (d) Fits of the chemical shift perturbations as function of C4 concentration (panel c).

To rationalize the observed increased affinity of C4A relative to C4, we modelled the pre-miR-31 complex both with C4 and C4A guided by the NMR HSQC data using Boltz-1.^34^ The modelled complex for the pre-miR-31-C4 and pre-miR-31-C4A interactions revealed similar binding poses. In both complexes the compounds form planar structures that stack between G29, U30, C42 and C43 at the dicing site. The presence of the OH group in C4A in place of the NH2 in C4 maintains similar hydrogen bonding interactions with the residue C42. The NH of the purine-like ring forms a hydrogen bonding interaction with G32. G29 forms hydrogen bonds with the two nitrogen atoms in the purine-like ring of both C4 and C4A. The presence of an extra NH2 group on the purine-like ring of C4A forms additional hydrogen bonding interactions with U30, increasing the overall interactions. The extra NH2 on the phenol of C4A may be solvent exposed since we do not observe any interactions with pre-miR-31 (**Fig. 7a, b**). While we do not have a structure of the RNA:ligand complexes, the additional interactions predicted for C4A are consistent with the substantially enhanced affinity of C4A for pre-miR-31 relative to C4.

**Figure 7:**
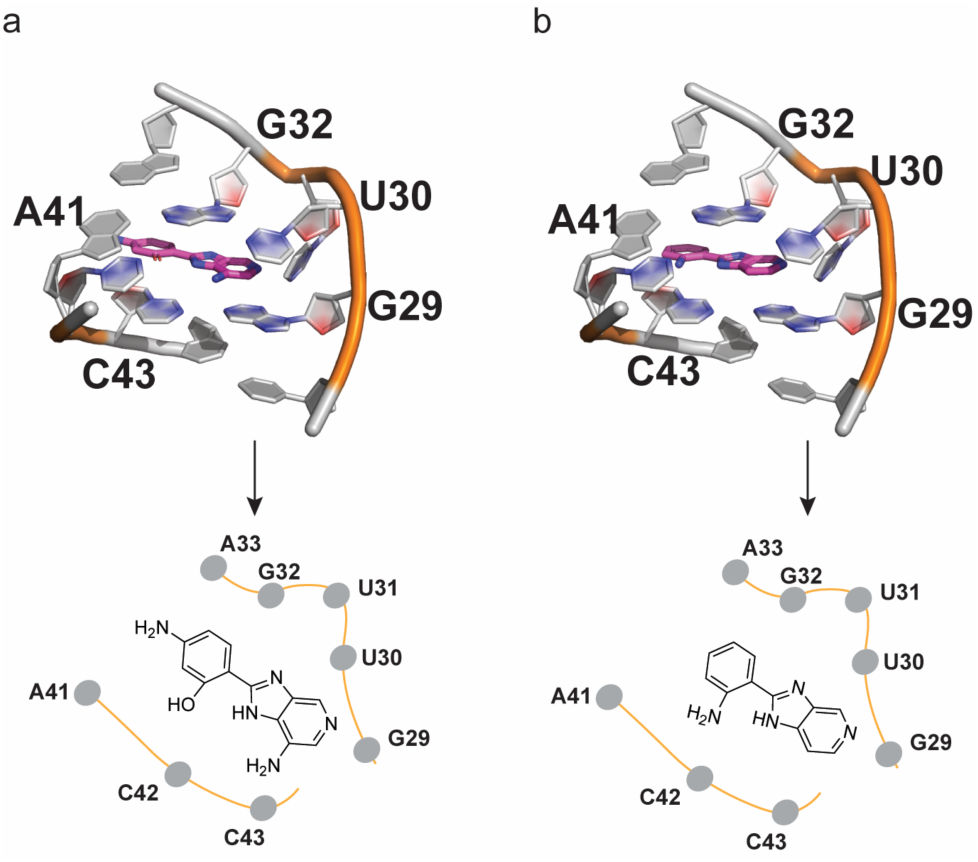
C4A and C4 bind at pre-miR-31 at the dicing loop. (a-b) Detailed view of (b) C4A and (c) C4 docked in pre-miR-31.

For full detailed comparisons we have included the predicted binding scores and apparent affinities of our compounds in **Table 1**.

**Table 1:** Predicted binding energies and apparent affinities of several confirmed hits.

| Small molecule | Predicted binding score(kcal/mol) | Apparent affinity, K<sub>D</sub> (μM)<sup>a</sup> |
| --- | --- | --- |
| DA1 | -19.893 | 20.64 ± 9.40 |
| C4A | -26.043 | 29.10 ± 0.15 |
| C4 | -18.690 | >1000 |
| A1 | -21.117 | >1000 |
| A1A | -21.355 | ND |
| A1B | -20.852 | ND |

<sup>a</sup> ND denotes affinities that were not detected due to no observable changes in the miR-31 hairpin spectrum at 10-fold excess of ligand

## Methods

### Virtual screening

We carried out virtual screening simulations on pre-miR-31 using a library of compounds (Diversity Set IV Library, Div4) curated by the National Cancer Institute (NCI) Developmental Therapeutics Program (DTP) containing 1,596 compounds. First, we obtained the 3D coordinates of the pre-miR-31 ensemble from the protein data bank (PDB ID: 8FCS).^20^ We screened for ligandable cavities within the ensemble using RNACavityMiner.^35^ Next, we docked Div4 compounds into the cavities identified within the pre-miR-31 ensemble using the rDock^36^ molecular docking software using our previously modified virtual screening protocol with few modifications.^24^ Docking was performed allowing 25 possible poses in the initial step. Hits were identified as compounds that were two standard deviations lower than the mean binding energy (most energetically favored). We used the miR-21 and miR-20b hairpins as selectivity controls by docking Div4 against these ensembles and eliminating compounds in our hit library that also bound these RNAs. Ultimately, we were left with 53 initial ‘hits’ and acquired top (highly scored) 40 compounds from the NCI for experimental characterization.

### DNA template preparation

We purchased the DNA template for the miR-31 hairpin construct from Integrated DNA Technologies (5ʹ-mGmGCATAGCAGGTTCCCAGTTCAACAGCTATGCCTATAGTGAGTCGTATTA-3ʹ). The template was 2ʹ-*O*-methoxy modified at the two 5ʹ most residues to reduce N + 1 product formation by T7 RNA polymerase.^37^ We prepared a partial DNA duplex required for transcription by annealing the template with Top-17; a 17-nucleotide DNA corresponding to the T7 promoter sequence (5ʹ-TAATACGACTCACTATA-3ʹ).

### RNA transcription

RNA was prepared by *in vitro* transcription with T7 RNA polymerase as previously described.^38^ Briefly, DNA template, ribonucleoside triphosphates (rNTPs), magnesium chloride (MgCl2), yeast inorganic pyrophosphatase (New England Biolabs), and dimethyl sulfoxide (DMSO)^39^ were mixed with in house prepared T7 RNA polymerase. The transcription reaction was incubated at 37 °C with shaking at 75 rpm. After 3 h, the reaction was quenched with a solution of 7 M urea and 500 mM ethylenediaminetetraacetic acid (EDTA) pH 8.5. RNA was purified from the crude transcription reaction by denaturing polyacrylamide gel electrophoresis. The quenched transcription reaction was loaded onto a 16% large scale denaturing polyacrylamide gel with the addition of 16% glycerol (50% v/v) and run for ∼15 h at 25 w. The RNA product was visualized with UV shadowing, excised from the gel, and extracted by electroelution. The eluted RNA was spin concentrated, washed with ultra-pure sodium chloride, and exchanged into water using a 3K Amicon Ultra centrifugal filter. RNA purity was confirmed on a 10% analytical denaturing polyacrylamide gel and the concentration was quantified via UV-Vis absorbance. ^15^N/^13^C-adenosine/cytosine (^15^N/^13^C-A/C) labelled RNA for HSQC experiments was prepared as described above except that unlabelled rATP and rCTP were replaced with ^15^N/^13^C-rATP and ^15^N/^13^C-rCTP in the transcription reaction.

### NMR data acquisition, processing, and analysis

^1^H spectra were obtained for all 40 compounds acquired from the NCI DTP and select derivatives (eMolecules). Samples were prepared by dissolving ligand in NMR buffer [500 mM KH2PO4, pH 7.5 and 1 mM MgCl2] containing 2% DMSO, and 10% D2O in a 550 μL volume to ∼500 μM concentration. Samples were transferred into a 5 mm NMR tube and spectra was acquired on a 600 MHz Bruker ADVANCE NEO spectrometer at 25 °C and processed with MestreNova (MNova).^40^

Samples for STD experiments were prepared in NMR buffer containing 2% DMSO, 98% D2O in a 550 μL volume to a final 10 μM RNA and ∼200 μM ligand concentration. The standard Bruker pulse program (stddiffesgp.3) was used with changes in resonance frequencies. The off-resonance frequency was set to -40 ppm for all compounds while the on-resonance frequency was set to either 4.9 or 5.5 ppm, depending on the ligand ^1^H spectra. Samples were transferred into a 5 mm NMR tube and spectra was acquired on a 600 MHz Bruker ADVANCE NEO spectrometer at 25 °C. The dataset was loaded into MNova as a stack of the off-resonance and on-resonance spectra. After processing, the spectra were subtracted from each other using the arithmetic function in MNova to generate the difference spectra.

Samples for HSQC experiments were prepared in NMR buffer containing 2% DMSO, 10% D2O in a 150 μL volume to a final 100 μM ^13^C/^15^N-A/C labelled RNA and ∼1 mM ligand concentration. Samples were transferred into a 3 mm NMR tube and spectra were acquired on a 800 MHz Bruker ADVANCE NEO spectrometer at 37 °C. Spectra were processed with NMRFx^41^ and analyzed with NMRViewJ.^42^

Titration experiments were performed to determine KD values of ligands for the pre-miR-31 RNA. The different concentrations used for the different ligands is highlighted in **Fig. 6 and Figs. S39, S40** (ranging from 0 – 1,000 μM). For compounds C4A and DA1, which were in intermediate exchange with the binding site residues, the KD was estimated by evaluating the changes in peak volumes as a function of ligand concentration by measuring the fraction bound at different concentrations using **Equation 1**.

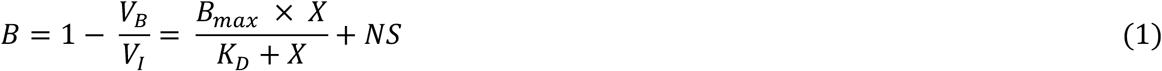

Where B is fraction bound at specific concentration, VB is the volume of the bound RNA peak and VI is the volume of unbound RNA peak, X is the total ligand concentration and NS is the slope of the nonlinear regression (nonspecific binding is assumed to be linear).

For C4 and A1, where addition of ligands was correlated to shifts in peak positions, we measured the chemical shift perturbation (CSP) values for peaks in the RNA using the Pythagorean distance moved in ^1^H and ^13^C dimensions, with weights attached to the ^13^C as shown in **Equation 2**.

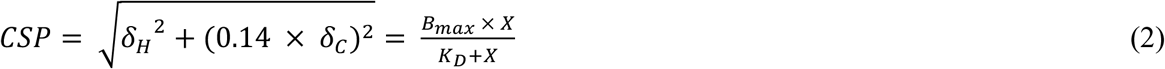

𝛿*_H_* is the distance moved in the proton dimension and 𝛿_*C*_ is the distance moved in the carbon dimension.

### Fluorescence indicator displacement assays

The affinity of TO-PRO-1 to the miR-31 hairpin was measured by varying the RNA concentration in 13 serial point dilutions (0 - 20 μM). Samples were prepared by dissolving the RNA into NMR buffer containing 2% DMSO on a clear 96-well plate. 50 μL of each dilution was transferred into a black costar 96 well plate. 50 μL of TO-PRO-1 (Invitrogen) prepared in NMR buffer containing 2% DMSO was then added to a final concentration of 500 nM to each well. The samples were incubated for 30 min at room temperature in the dark and the plates were read on a SpectraMax iD3 Plate Reader (excitation = 492 nm, absorption = 575 nm). Data were fit to a one site binding curve (**Equation 3**) using GraphPad Prism (GraphPad Software, La Jolla California USA, www.graphpad.com).

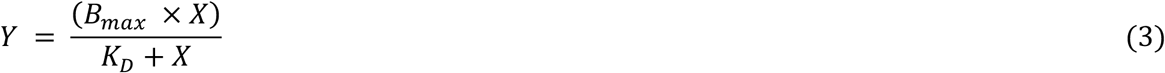

Where X is RNA concentration, Y is fluorescence intensity, and 𝐵_*max*_ is the highest fluorescence (RFU). The resulting KD was used to calculate the value of X (RNA concentration) at an initial fraction bound at 0.1.

For the screening assay, samples were prepared in a black costar 96 well plate. RNA-TO-PRO-1 complex was prepared at 90 μL to a final concentration 50.9 nM RNA and 500 nM TO-PRO-1 in 100 μL and incubated in the dark for 30 min. Following incubation, ligands dissolved in NMR buffer containing 2% DMSO were added in 10 μL volumes to a final concentration of 100 μM. Plates were shaken for 1 min at 600 rpm and centrifuged at 2510 x g for 1 min and incubated in the dark for 30 min and then read on a SpectraMax iD3 Plate Reader (excitation = 492 nm, absorption = 575 nm). Percent fluorescence indicator displacement (%FID) using **Equation 4**.

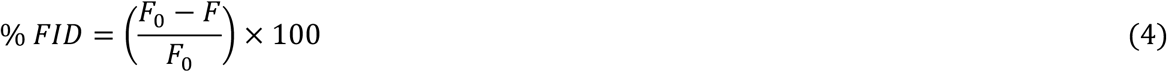

Where *F0* is the fluorescence of the blank well with RNA + dye and no small molecule and *F* is the fluorescence of the well with all three components (RNA + dye + small molecule). Changes in fluorescence used in the plot were achieved by subtracting the %FID from 100 (100-%FID). Screening was performed in duplicate. Note RNA-ligand controls as well as ligand-dye controls were performed and analyzed as described except in the absence of the dye or the RNA, respectively.

### Conclusions

We have described an integrative multi-screening protocol to discover small molecules that bind the miR-31 hairpin. We identified 40 initial ‘hits’ by virtually screening a library containing 1,596 compounds against ligandable cavities within pre-miR-31 and followed up with binding experiments using both STD-NMR and a fluorescence indicator displacement assay. Individually, these experiments have been successfully applied to screen for RNA-small molecule binders. We applied these approaches simultaneously to validate small molecule hits identified from our virtual screening campaign. Our STD-NMR resulted in an initial hit rate of ∼18% while our indicator displacement assay yielded a 10% hit rate. These differences highlighted the inherent limitations in both assays. STD-NMR is constrained by compounds solubility because of the requirement for higher sample concentration. In contrast, fluorescence indicator displacement assays can be influenced by the inherent fluorescent properties of small molecules upon their contact with an indicator. Using both methods simultaneously allowed us to gain a more complete understanding of the RNA:ligand interactions. This dual assay approach improved our success at small molecule discovery and collectively identified 25% of our compounds identified from the *in silico* screening are true binders of the miR-31 hairpin. This hit rate, while comparable to those reported in other virtual screening campaigns, highlights limitations of current scoring functions. This gap can only be bridged as more RNA-small molecule binding data becomes available to improve the predictive accuracy of current scoring functions. Strikingly, three compounds from our validated hits showed chemical shift perturbations in the miR-31 hairpin at the junction region and dicing site residues. To enrich our hit library further, we used two of our initial scaffolds as basis to chemical structure similarity search to identify additional ligands that bound the miR-31 hairpin. Among the newly discovered small molecules we found that C4A, a derivative of C4, bound the miR-31 hairpin with >20-fold affinity compared to C4. Furthermore, a docked model of the pre-miR-31-C4A and pre-miR-31-C4 complex revealed that both compounds bound the dicing site of the miR-31 hairpin.

Our study has offered critical insights into the miR-31 harpin-small molecule recognition, providing foundational knowledge on the ligandability of pre-miR-31. Notably the small molecules with the highest affinity for the miR-31 hairpin (C4A and DA1) bound with low micromolar affinities, affinities that are likely too weak to impact biological function. However, these binders serve as valuable starting points for further optimization including affinity improvements or the design of more sophisticated molecules such as proximity induced degraders and ribonuclease targeting chimeras (RIBOTACs) to achieve functionality.^43,44^

## ACKNOWLEDGEMENTS

This work was supported by National Institute of General Medical Sciences of the National Institutes of Health grant R35 GM138279 (to S.C.K.) and Research Corporation for Science Advancement Cottrell Scholar Award 28248 (S.C.K.). Research reported in this publication was supported by the University of Michigan BioNMR Core Facility (U-M BioNMR) and a research grant from the Rackham graduate school. Compounds used in the initial screening were provided by the National Cancer Institute (NCI) Developmental Therapeutics Program. We thank Prof. Charles L. Brooks III for giving us access to the Gollum computing cluster and Debashish Sahu and Minli Xing for technical assistance with NMR data acquisition. Finally, we thank Dr. Aaron Frank (Arrakis Therapeutics) for valuable discussions and useful contributions to the development of this project.

## AUTHOR CONTRIBUTIONS

G.A. and S.C.K. conceived the project. G.A. and L.H. performed and analyzed indicator displacement assays. G.A performed and analyzed all other work. S.C.K supervised the project. G.A and S.C.K. wrote the manuscript.

## DATA AVAILABILITY

The data supporting this article has been included as part of the Supplementary Information.

## References

1 W. Chauhan, Sudharshan Sj, S. Kafle and R. Zennadi, Int J Mol Sci, 2024, 25, 7202.

2 D. P. Bartel, Cell, 2004, 116, 281–297.

3 R. Bayraktar and K. Van Roosbroeck, Cancer Metastasis Rev, 2018, 37, 33–44.

4 J. L. Childs-Disney, X. Yang, Q. M. R. Gibaut, Y. Tong, R. T. Batey and M. D. Disney, Nat Rev Drug Discov, 2022, 21, 736–762.

5 S. P. Velagapudi, M. D. Cameron, C. L. Haga, L. H. Rosenberg, M. Lafitte, D. R. Duckett, D. G. Phinney and M. D. Disney, Proc. Natl. Acad. Sci. U.S.A., 2016, 113, 5898–5903.

6 Y. Du, Y. Lin, K. Yin, L. Zhou, Y. Jiang, W. Yin and J. Lu, Am J Transl Res, 2019, 11, 1748– 1759.

7 J. Hayes, P. P. Peruzzi and S. Lawler, Trends in Molecular Medicine, 2014, 20, 460–469.

8 T. Kawamata and Y. Tomari, Trends Biochem Sci, 2010, 35, 368–376.

9 E. Huntzinger and E. Izaurralde, Nat Rev Genet, 2011, 12, 99–110.

10 Y. Lee, M. Kim, J. Han, K.-H. Yeom, S. Lee, S. H. Baek and V. N. Kim, EMBO J, 2004, 23, 4051–4060.

11 J. Han, Y. Lee, K.-H. Yeom, Y.-K. Kim, H. Jin and V. N. Kim, Genes Dev., 2004, 18, 3016– 3027.

12 J. Han, Y. Lee, K.-H. Yeom, J.-W. Nam, I. Heo, J.-K. Rhee, S. Y. Sohn, Y. Cho, B.-T. Zhang and V. N. Kim, Cell, 2006, 125, 887–901.

13 S. C. Kwon, T. A. Nguyen, Y.-G. Choi, M. H. Jo, S. Hohng, V. N. Kim and J.-S. Woo, Cell, 2016, 164, 81–90.

14 Y. Lee, C. Ahn, J. Han, H. Choi, J. Kim, J. Yim, J. Lee, P. Provost, O. Rådmark, S. Kim and V. N. Kim, Nature, 2003, 425, 415–419.

15 M. Fareh, K.-H. Yeom, A. C. Haagsma, S. Chauhan, I. Heo and C. Joo, Nat Commun, 2016, 7, 13694.

16 R. F. Ketting, S. E. J. Fischer, E. Bernstein, T. Sijen, G. J. Hannon and R. H. A. Plasterk, Genes Dev., 2001, 15, 2654–2659.

17 Z. Liu, J. Wang, H. Cheng, X. Ke, L. Sun, Q. C. Zhang and H.-W. Wang, Cell, 2018, 173, 1191–1203.e12.

18 G. Michlewski and J. F. Cáceres, RNA, 2019, 25, 1–16.

19 B. Wightman, I. Ha and G. Ruvkun, Cell, 1993, 75, 855–862.

20 S. Ma, A. Kotar, I. Hall, S. Grote, S. Rouskin and S. C. Keane, Proc. Natl. Acad. Sci. U.S.A., 2023, 120, e2300527120.

21 S. Ma, S. A. Howden and S. C. Keane, bioRxiv, 2024, 2024.04.08.588531.

22 O. Slaby, M. Svoboda, P. Fabian, T. Smerdova, D. Knoflickova, M. Bednarikova, R. Nenutil and R. Vyzula, Oncology, 2007, 72, 397–402.

23 N. Wang, Y. Li and J. Zhou, BioMed Research International, 2017, 2017, 1–12.

24 G. Arhin and S. C. Keane, Biophysics, 2025, preprint, DOI: 10.1101/2025.05.07.652683.

25 A. Viegas, J. Manso, F. L. Nobrega and E. J. Cabrita, J. Chem. Educ., 2011, 88, 990–994.

26 M. Mayer and T. L. James, J Am Chem Soc, 2004, 126, 4453–4460.

27 I. Gómez Pinto, C. Guilbert, N. B. Ulyanov, J. Stearns and T. L. James, J. Med. Chem., 2008, 51, 7205–7215.

28 L. T. Olenginski, W. K. Kasprzak, S. K. Attionu, B. A. Shapiro and T. K. Dayie, Molecules, 2023, 28, 1803.

29 M. Zafferani, J. G. Martyr, D. Muralidharan, N. I. Montalvan, Z. Cai and A. E. Hargrove, ACS Chem. Biol., 2022, 17, 2437–2447.

30 R. Del Villar-Guerra, R. D. Gray, J. O. Trent and J. B. Chaires, Nucleic Acids Res, 2018, 46, e41.

31 D. L. Boger, B. E. Fink, S. R. Brunette, W. C. Tse and M. P. Hedrick, J Am Chem Soc, 2001, 123, 5878–5891.

32 D. R. Calabrese, C. M. Connelly and J. S. Schneekloth, in Methods in Enzymology, Elsevier, 2019, vol. 623, pp. 131–149.

33 A. Cereto-Massagué, M. J. Ojeda, C. Valls, M. Mulero, S. Garcia-Vallvé and G. Pujadas, Methods, 2015, 71, 58–63.

34 J. Wohlwend, G. Corso, S. Passaro, N. Getz, M. Reveiz, K. Leidal, W. Swiderski, L. Atkinson, T. Portnoi, I. Chinn, J. Silterra, T. Jaakkola and R. Barzilay, Cold Spring Harbor Laboratory, 2024, preprint, DOI: 10.1101/2024.11.19.624167.

35 J. Xie and A. T. Frank, ACS Med Chem Lett, 2021, 12, 928–934.

36 S. Ruiz-Carmona, D. Alvarez-Garcia, N. Foloppe, A. B. Garmendia-Doval, S. Juhos, P. Schmidtke, X. Barril, R. E. Hubbard and S. D. Morley, PLoS Comput Biol, 2014, 10, e1003571.

37 C. Kao, M. Zheng and S. Rüdisser, RNA, 1999, 5, 1268–1272.

38 Y. Liu, A. Munsayac, I. Hall and S. C. Keane, J Mol Biol, 2022, 167688.

39 P. G. Popova, M. A. Lagace, G. Tang and A. K. Blakney, Eur J Pharm Biopharm, 2024, 198, 114247.

40 M. R. Willcott, J. Am. Chem. Soc., 2009, 131, 13180–13180.

41 M. Norris, B. Fetler, J. Marchant and B. A. Johnson, J Biomol NMR, 2016, 65, 205–216.

42 B. A. Johnson, Methods Mol Biol, 2004, 278, 313–352.

43 M. G. Costales, Y. Matsumoto, S. P. Velagapudi and M. D. Disney, J. Am. Chem. Soc., 2018, 140, 6741–6744.

44 S. Mikutis, M. Rebelo, E. Yankova, M. Gu, C. Tang, A. R. Coelho, M. Yang, M. E. Hazemi, M. Pires De Miranda, M. Eleftheriou, M. Robertson, G. S. Vassiliou, D. J. Adams, J. P. Simas, F. Corzana, J. S. Schneekloth, K. Tzelepis and G. J. L. Bernardes, ACS Cent. Sci., 2023, 9, 892– 904.

