## Supplementary Data for "Integrated *in silico* and experimental screening identifies novel ligands that target precursor microRNA-31 at the Dicer cleavage site"

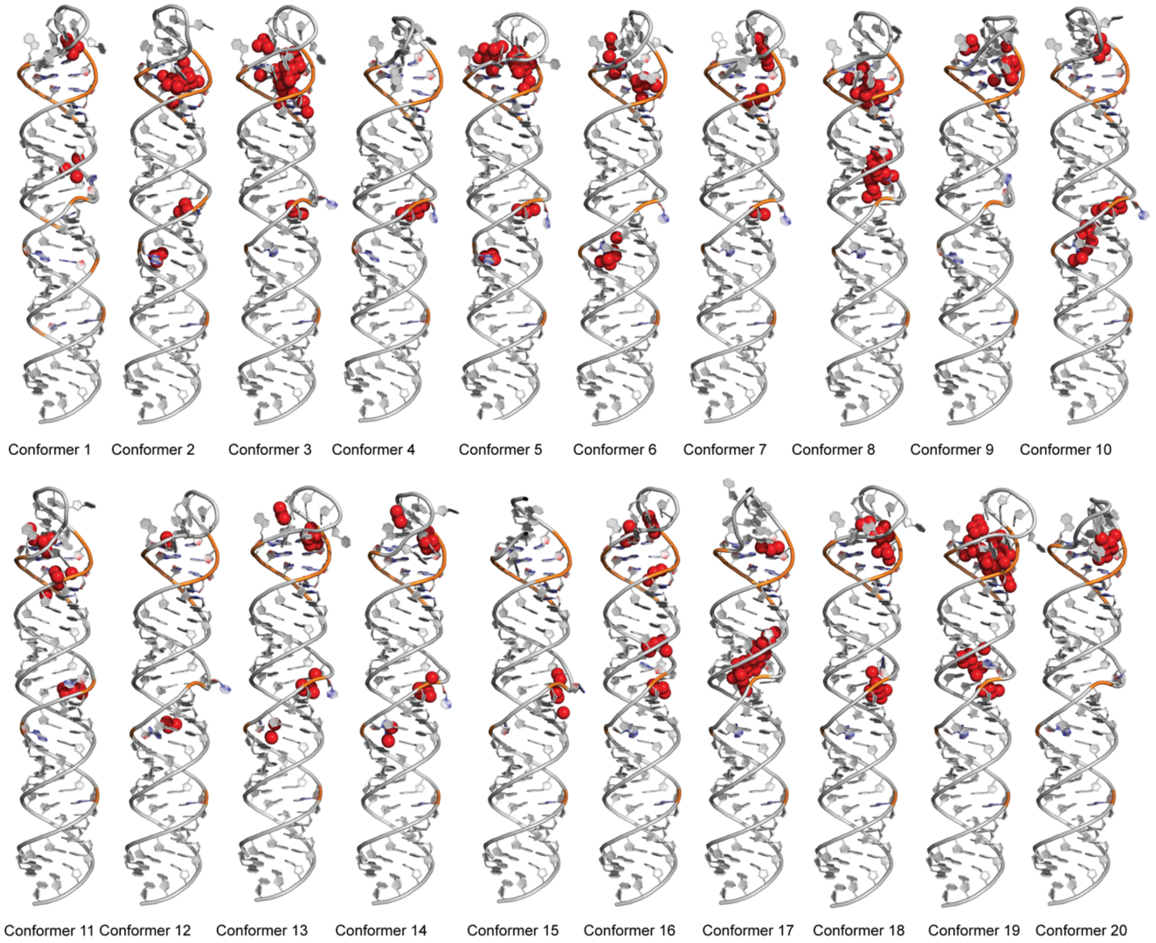


**Figure S1.** 3D structure of pre-miR-31 conformer showing predicted binding cavities (highlighted with placeholder atoms in red) determined using RNACavityMiner.

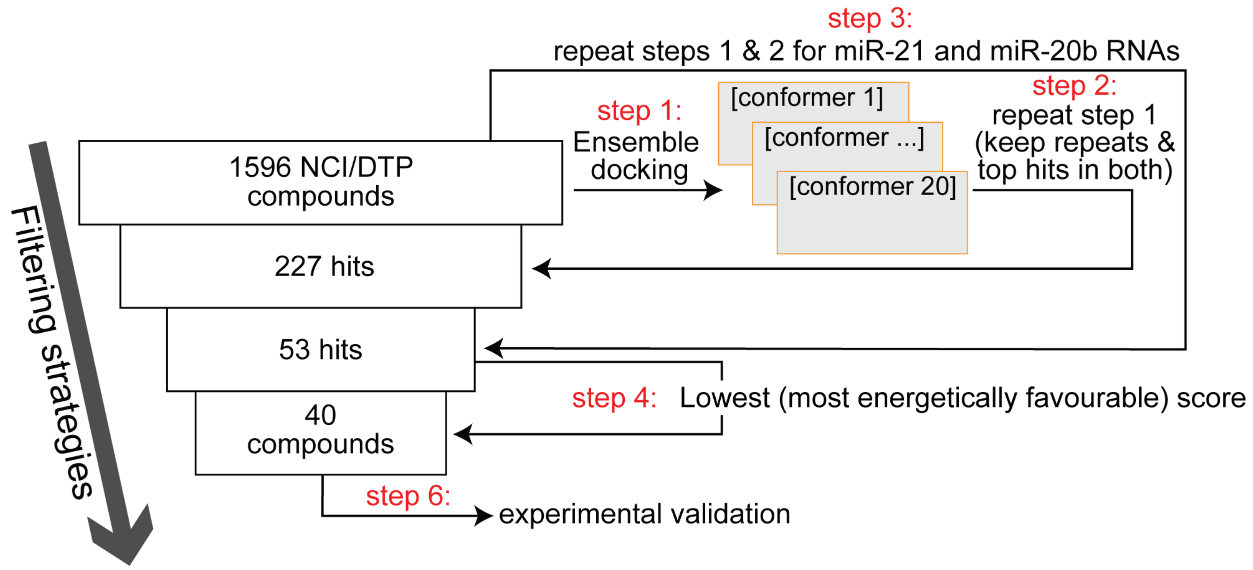


**Figure S2.** Schematic of the ensemble based virtual screening pipeline.

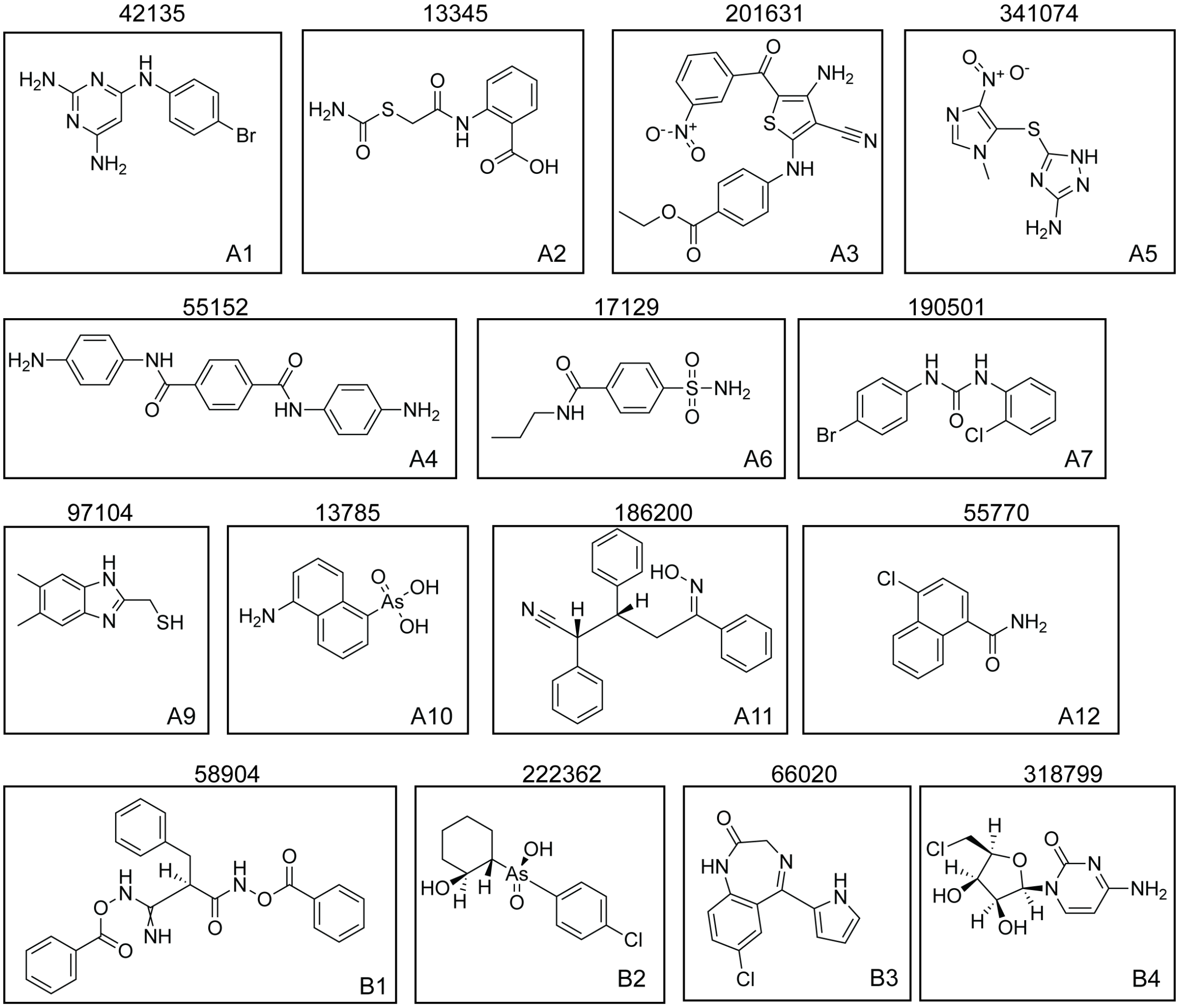


**Figure S3.** Chemical structures of compounds acquired from NCI indicating NCI ID number (top) and new assigned name (bottom right).

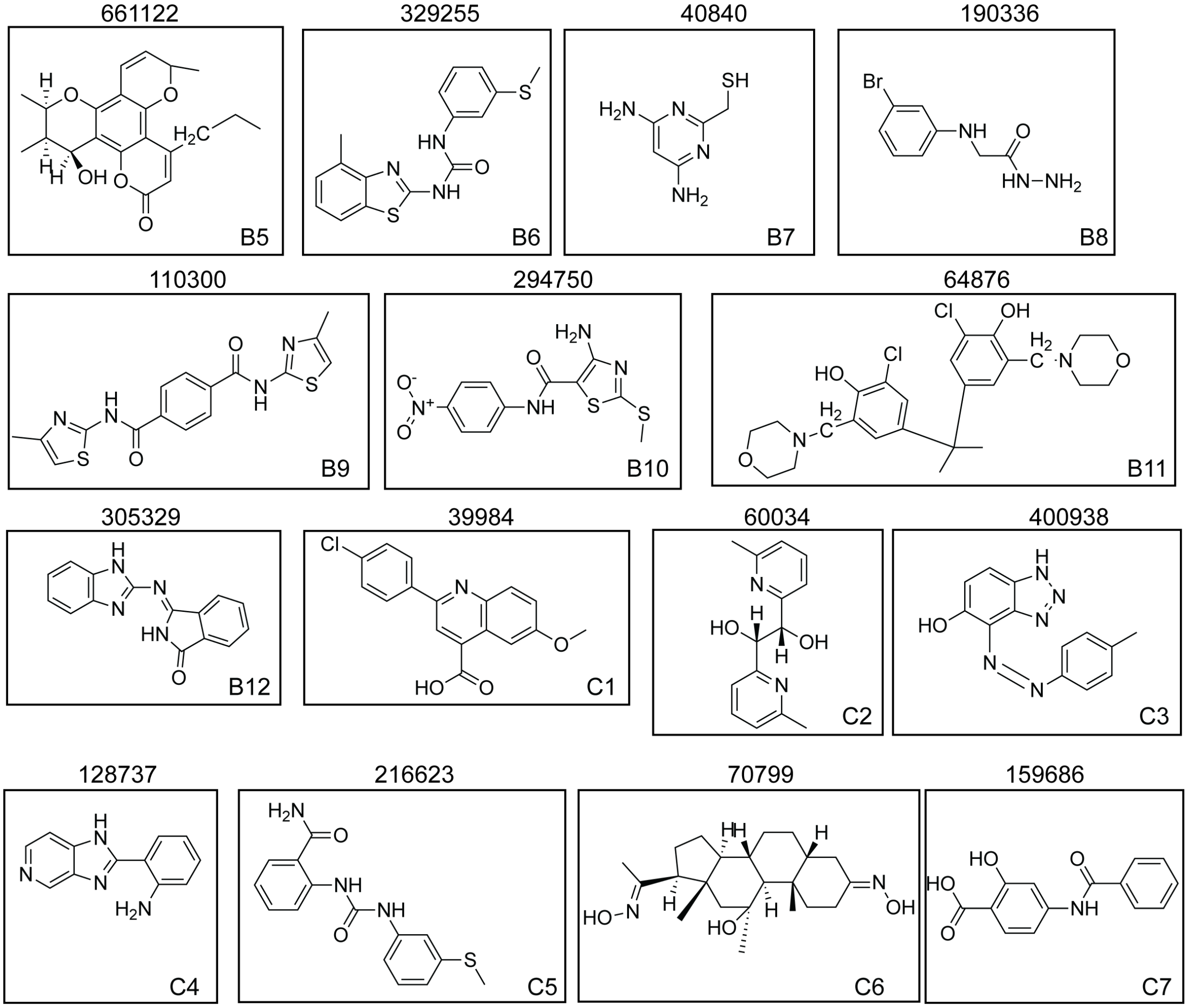


**Figure S4.** Chemical structures of compounds acquired from NCI indicating NCI ID number (top) and new assigned name (bottom right).

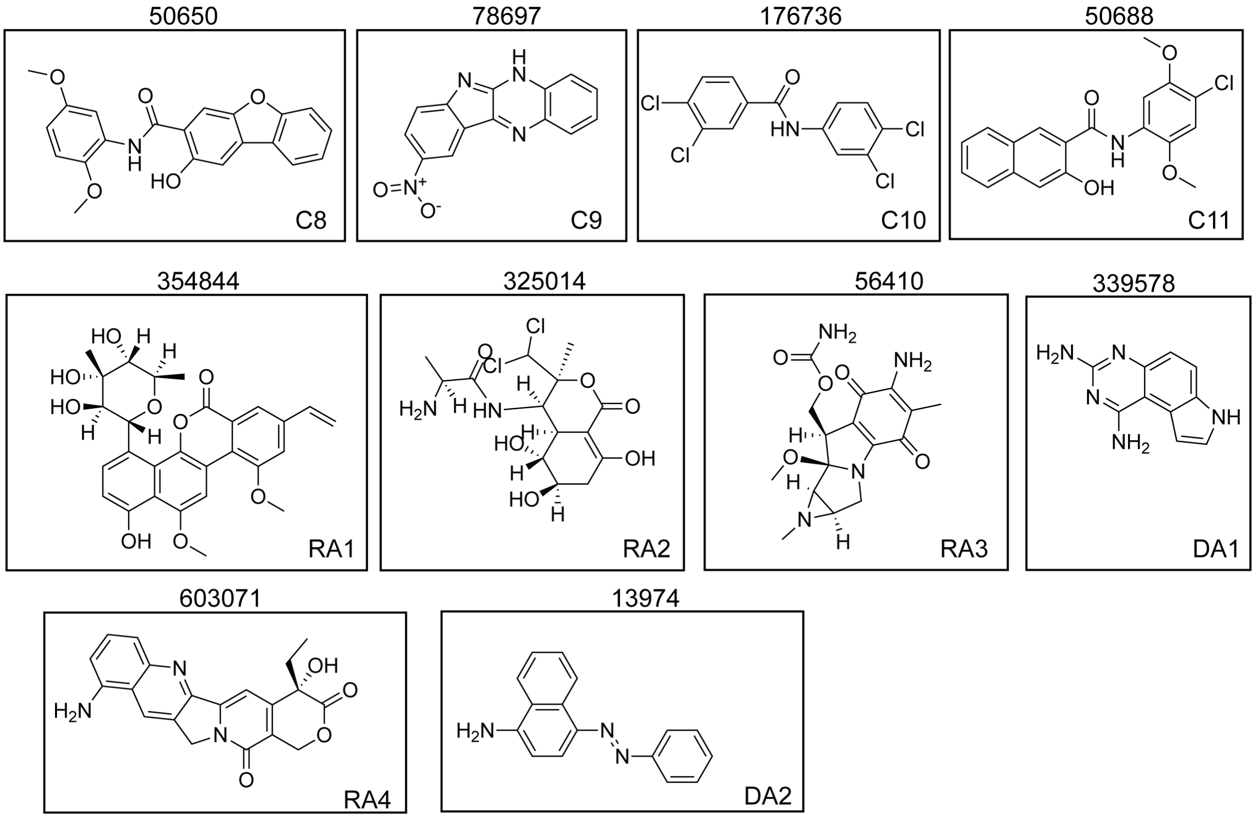


**Figure S5.** Chemical structures of compounds acquired from NCI indicating NCI ID number (top) and new assigned name (bottom right).

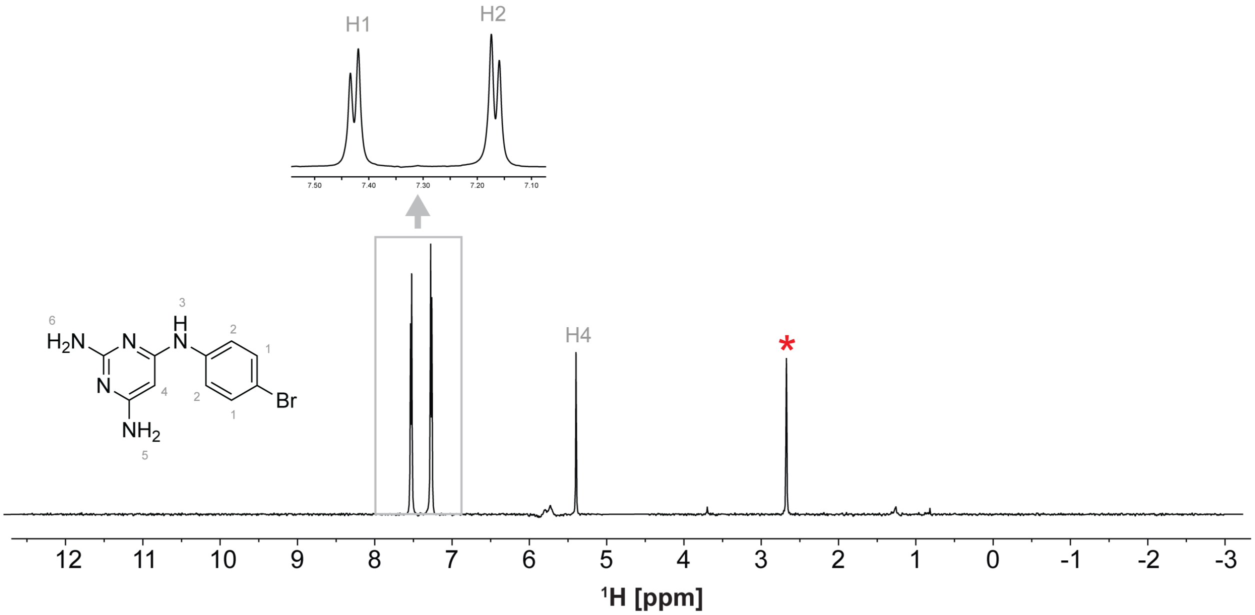


**Figure S6.** Chemical structure and $^1$H spectrum of A1. The NMR spectrum was recorded at ~ 500 µM concentration, 50 mM $KH_2PO_4$, pH = 7.5, 50 mM KCl, 1 mM $MgCl_2$, 10% $D_2O$ and 2% DMSO on a 600 MHz Bruker spectrometer equipped with a cryoprobe. The red asterisk indicates the DMSO peak.

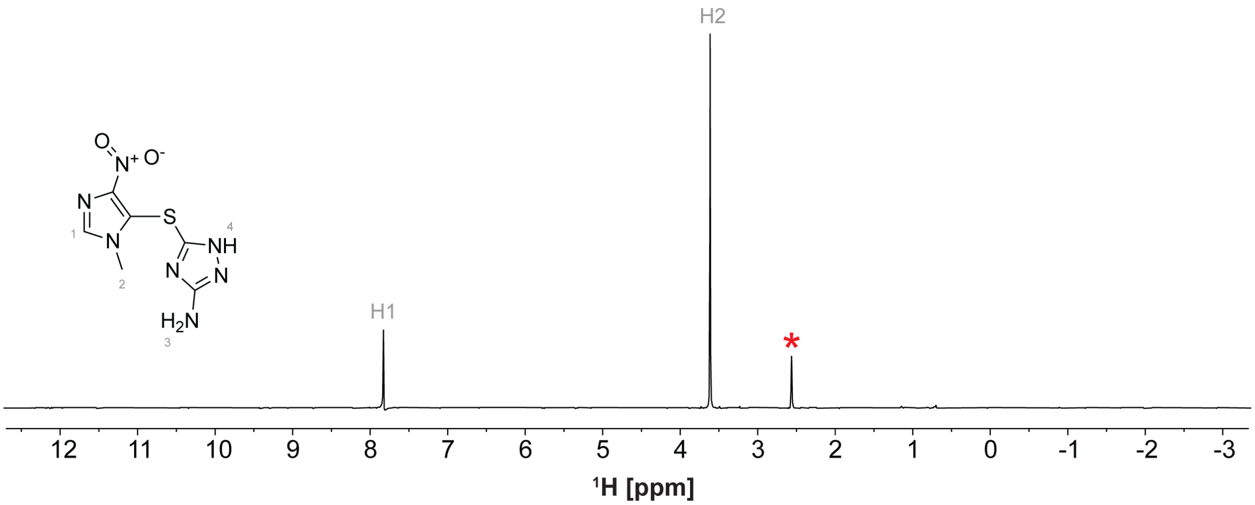


**Figure S7.** Chemical structure and $^1$H spectrum of A5. The NMR spectrum was recorded at ~ 500 μM concentration, 50 mM $KH_2PO_4$, pH = 7.5, 50 mM KCl, 1 mM $MgCl_2$, 10% $D_2O$ and 2% DMSO on a 600 MHz Bruker spectrometer equipped with a cryoprobe. The red asterisk indicates the DMSO peak.

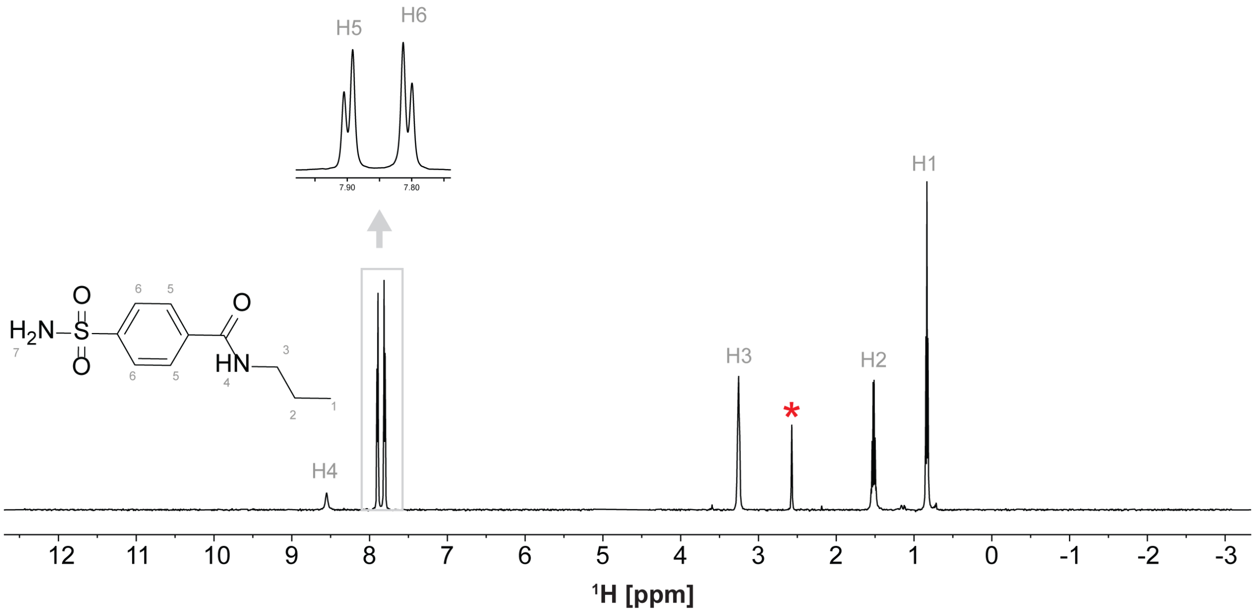


**Figure S8.** Chemical structure and [1]H spectrum of A6. The NMR spectrum was recorded at ~ 500 μM concentration, 50 mM $KH_2PO_4$, pH = 7.5, 50 mM KCl, 1 mM $MgCl_2$, 10% $D_2O$ and 2% DMSO on a 600 MHz Bruker spectrometer equipped with a cryoprobe. The red asterisk indicates the DMSO peak.

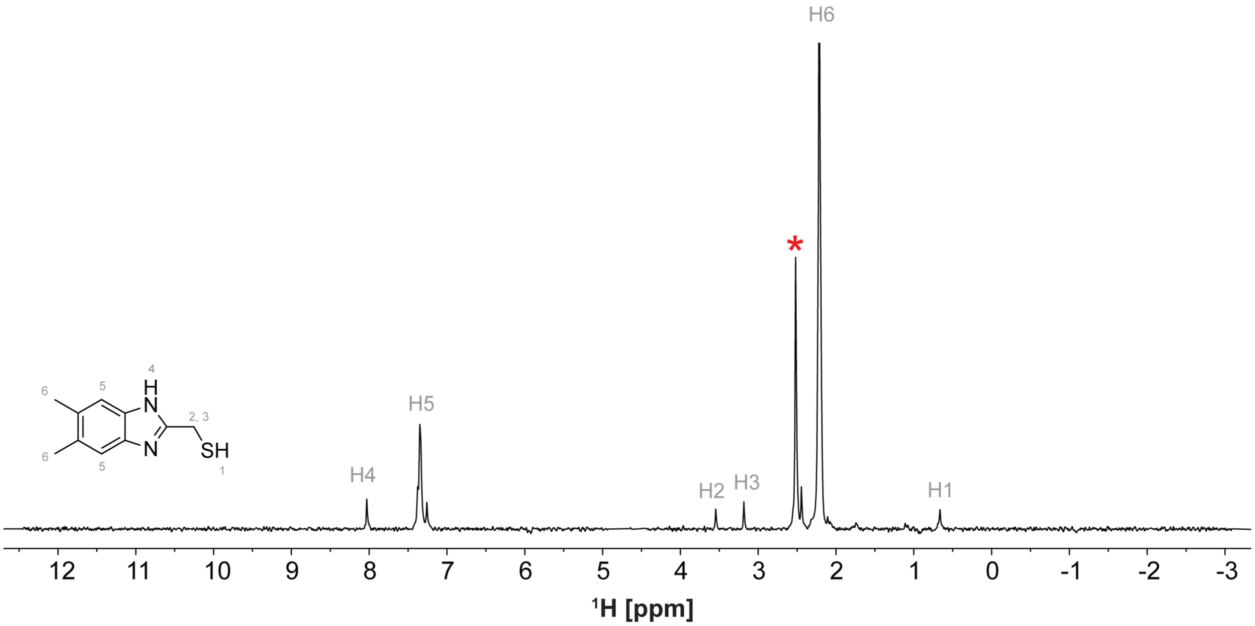


**Figure S9.** Chemical structure and ¹H spectrum of A9. The NMR spectrum was recorded at ~ 500 µM concentration, 50 mM $KH_2PO_4$, pH = 7.5, 50 mM KCl, 1 mM $MgCl_2$, 10% $D_2O$ and 2% DMSO on a 600 MHz Bruker spectrometer equipped with a cryoprobe. The red asterisk indicates the DMSO peak.

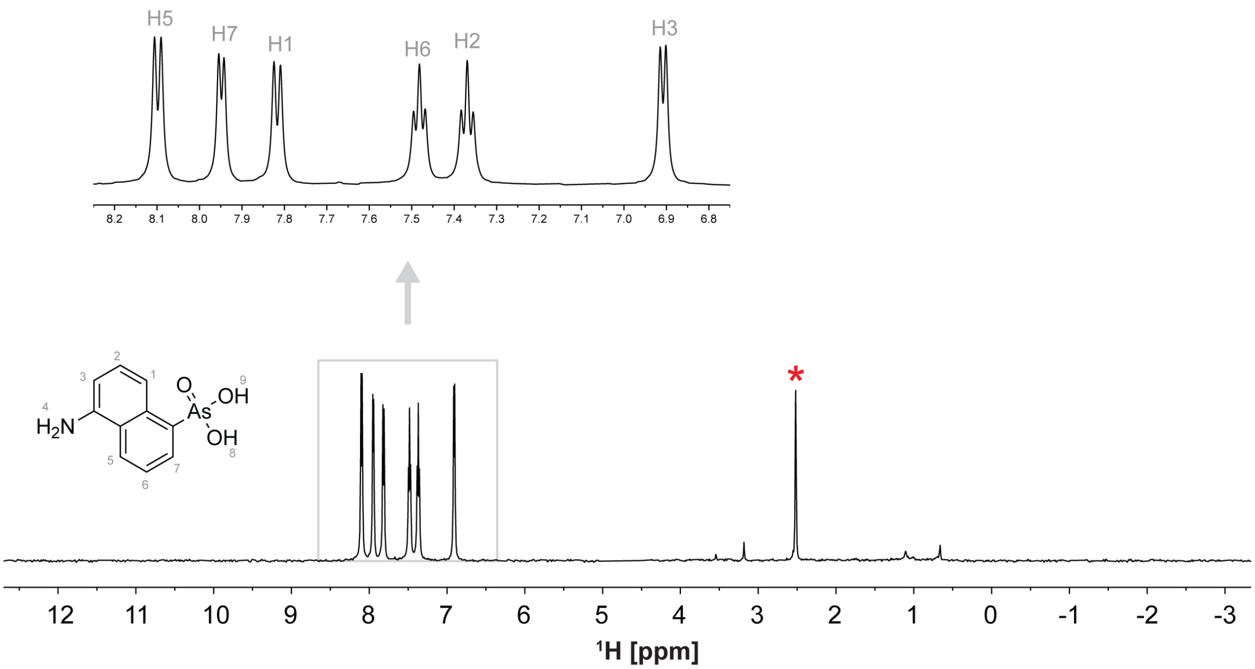


**Figure S10.** Chemical structure and $^{1}H$ spectrum of A10. The NMR spectrum was recorded at ~ 500 µM concentration, 50 mM $KH_2PO_4$, pH = 7.5, 50 mM KCl, 1 mM $MgCl_2$, 10% $D_2O$ and 2% DMSO on a 600 MHz Bruker spectrometer equipped with a cryoprobe. The red asterisk indicates the DMSO peak.

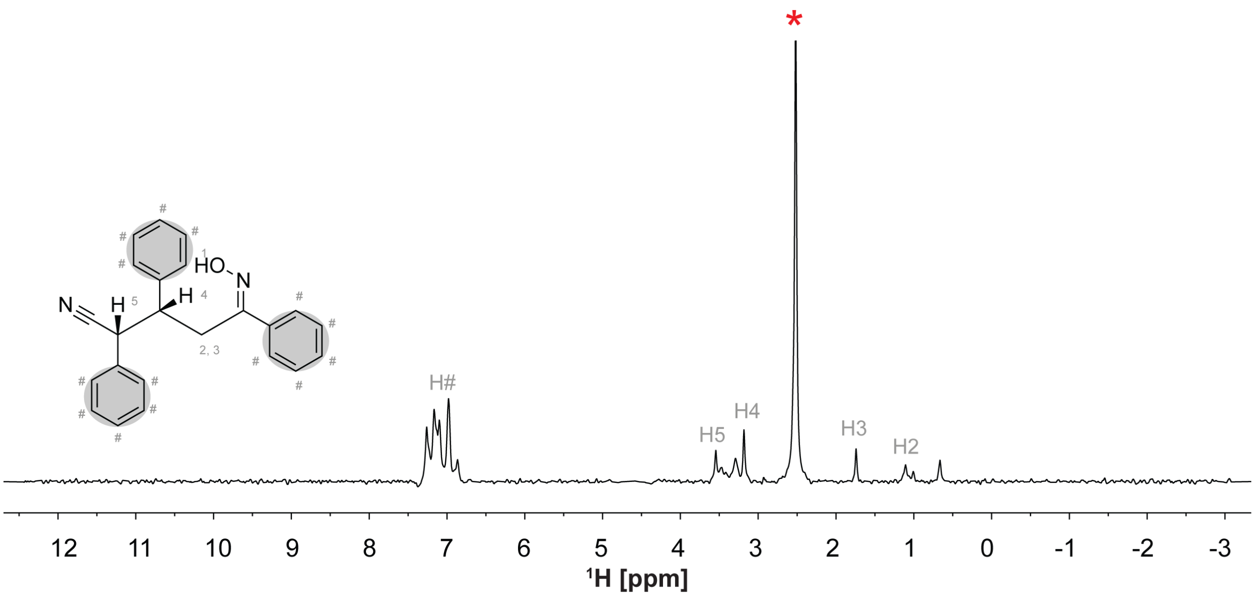


**Figure S11.** Chemical structure and $^{1}$H spectrum of A11. The NMR spectrum was recorded at ~ 500 µM concentration, 50 mM $KH_2PO_4$, pH = 7.5, 50 mM KCl, 1 mM $MgCl_2$, 10% $D_2O$ and 2% DMSO on a 600 MHz Bruker spectrometer equipped with a cryoprobe. The red asterisk indicates the DMSO peak.

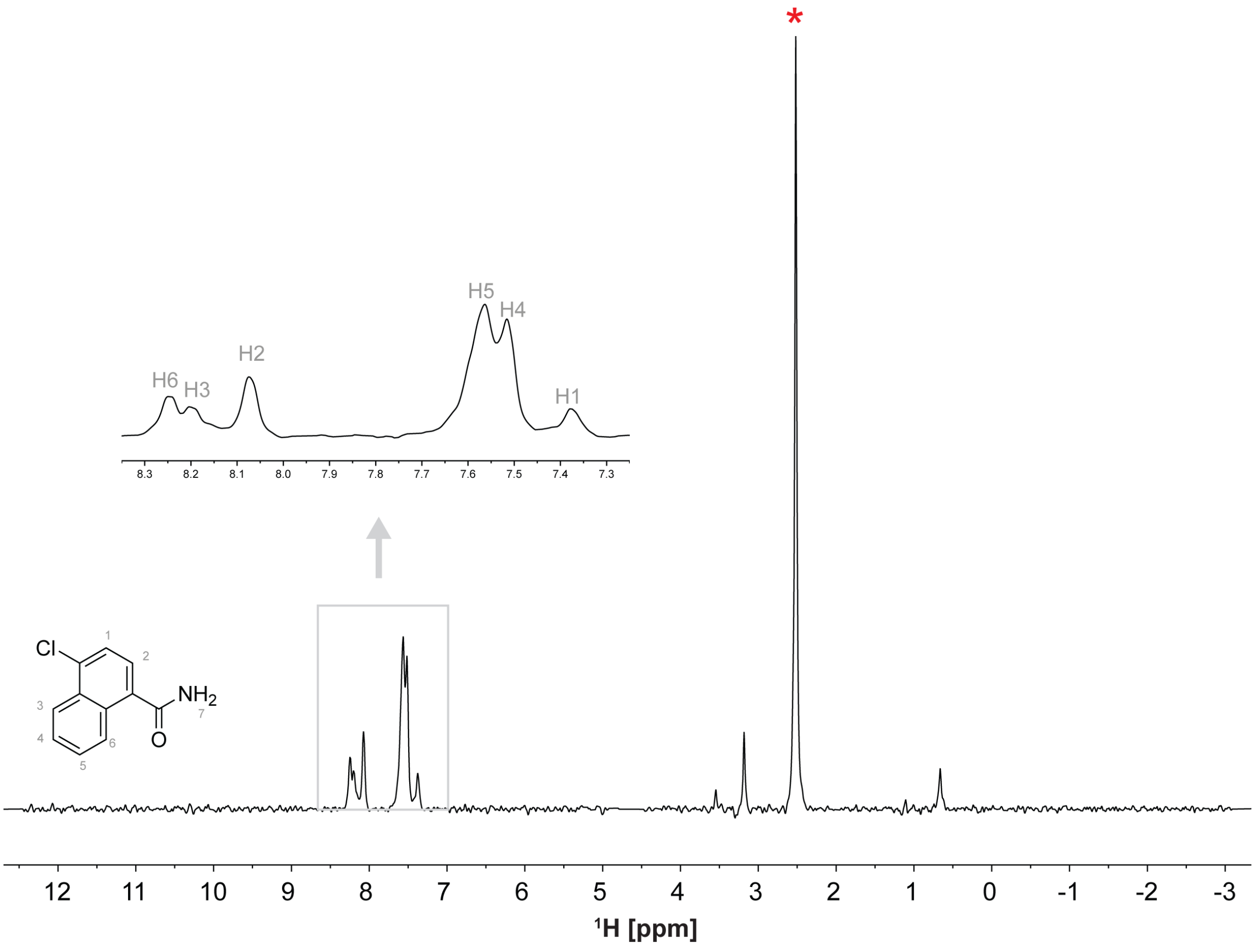


**Figure S12.** Chemical structure and $^1H$ spectrum of A12. The NMR spectrum was recorded at ~ 500 µM concentration, 50 mM $KH_2PO_4$, pH = 7.5, 50 mM KCl, 1 mM $MgCl_2$, 10% $D_2O$ and 2% DMSO on a 600 MHz Bruker spectrometer equipped with a cryoprobe. The red asterisk indicates the DMSO peak.

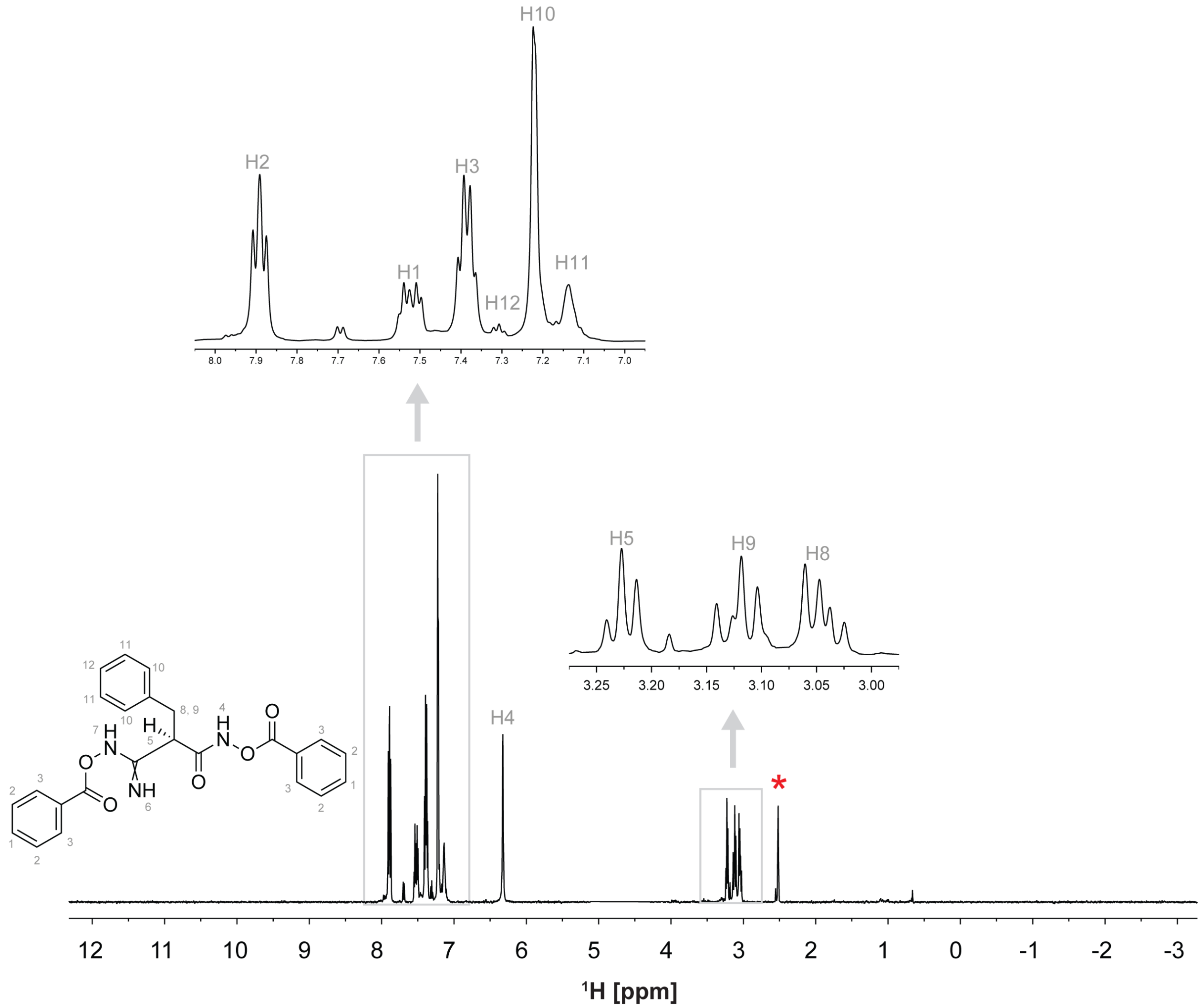


**Figure S13.** Chemical structure and $^{1}H$ spectrum of B1. The NMR spectrum was recorded at ~ 500 µM concentration, 50 mM $KH_2PO_4$, pH = 7.5, 50 mM KCl, 1 mM $MgCl_2$, 10% $D_2O$ and 2% DMSO on a 600 MHz Bruker spectrometer equipped with a cryoprobe. The red asterisk indicates the DMSO peak.

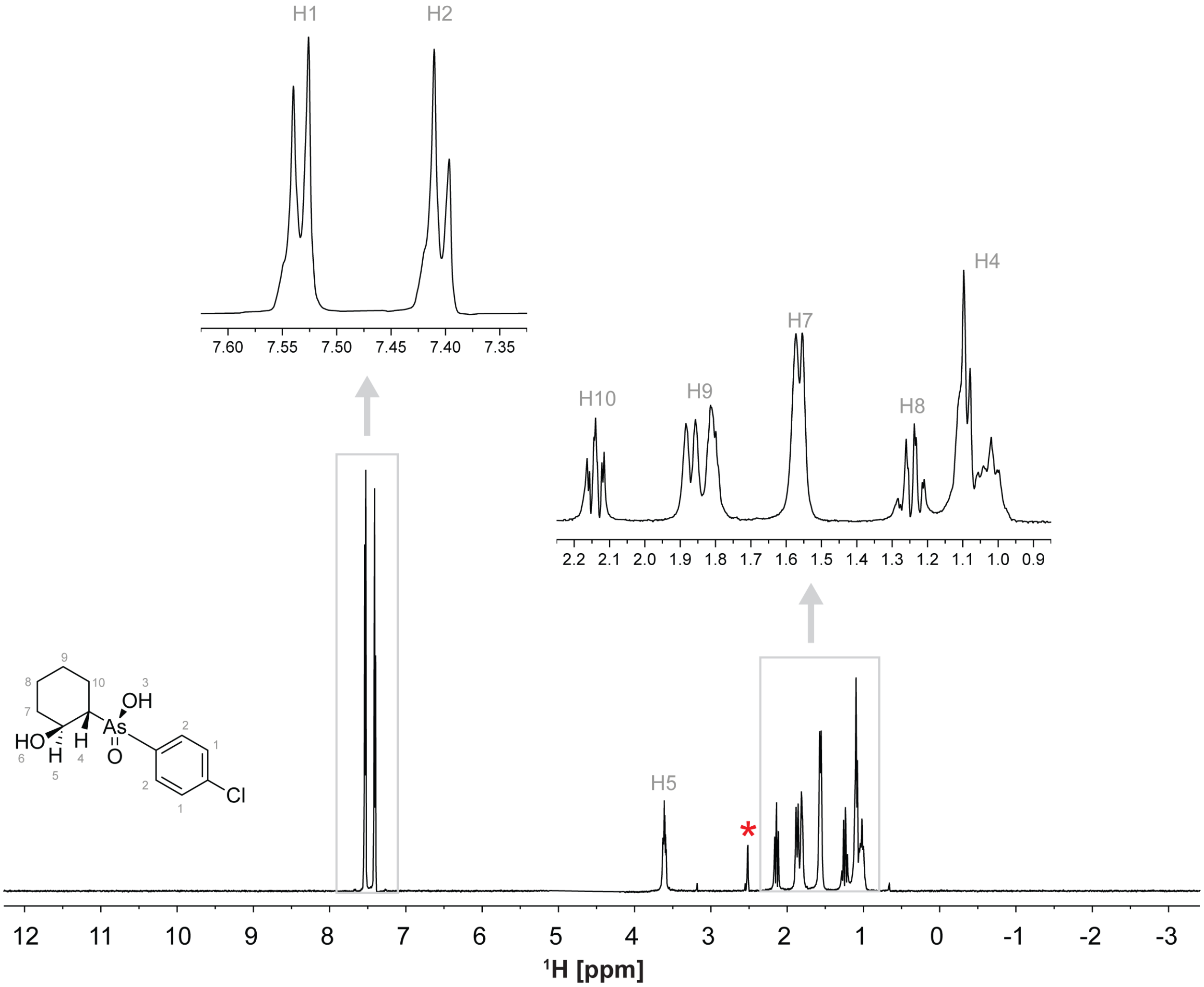


**Figure S14.** Chemical structure and $^1$H spectrum of B2. The NMR spectrum was recorded at ~ 500 µM concentration, 50 mM $KH_2PO_4$, pH = 7.5, 50 mM KCl, 1 mM $MgCl_2$, 10% $D_2O$ and 2% DMSO on a 600 MHz Bruker spectrometer equipped with a cryoprobe. The red asterisk indicates the DMSO peak.

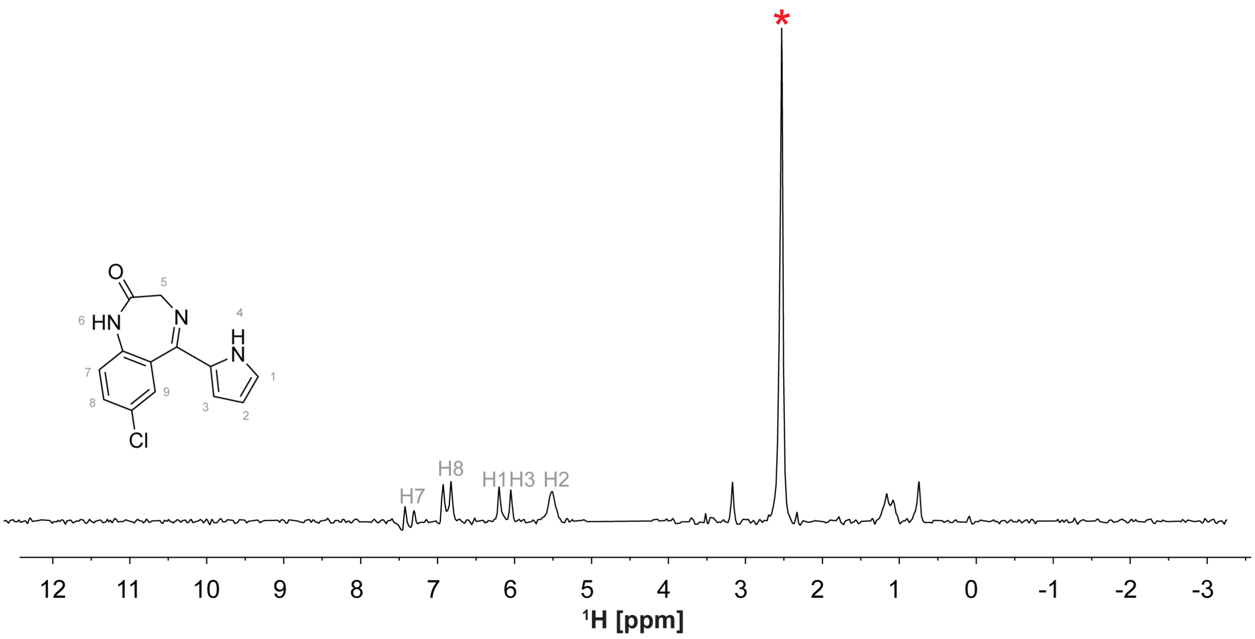


**Figure S15.** Chemical structure and [1]H spectrum of B3. The NMR spectrum was recorded at ~ 500 µM concentration, 50 mM $KH_2PO_4$, pH = 7.5, 50 mM KCl, 1 mM $MgCl_2$, 10% $D_2O$ and 2% DMSO on a 600 MHz Bruker spectrometer equipped with a cryoprobe. The red asterisk indicates the DMSO peak.

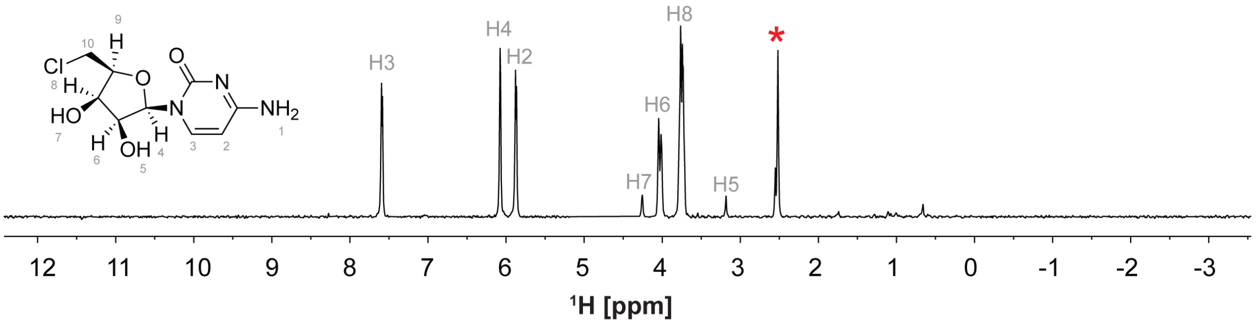


**Figure S16.** Chemical structure and $^1$H spectrum of B4. The NMR spectrum was recorded at ~ 500 µM concentration, 50 mM $KH_2PO_4$, pH = 7.5, 50 mM KCl, 1 mM $MgCl_2$, 10% $D_2O$ and 2% DMSO on a 600 MHz Bruker spectrometer equipped with a cryoprobe. The red asterisk indicates the DMSO peak.

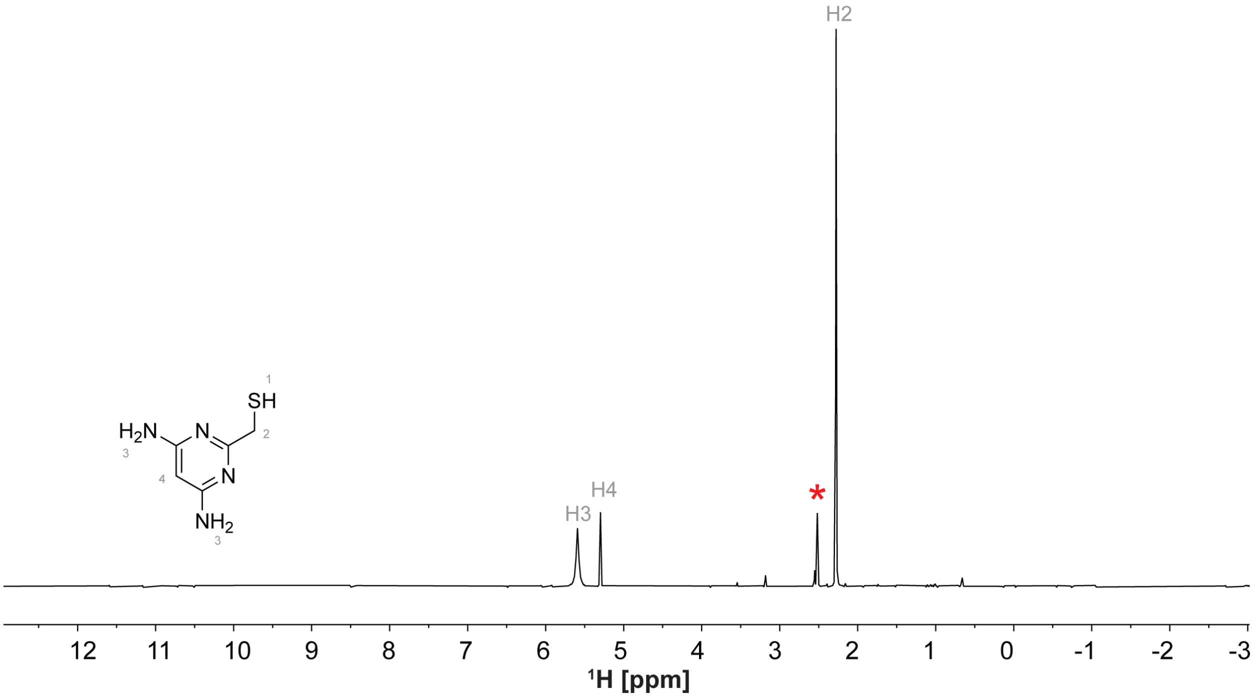


**Figure S17.** Chemical structure and $^{1}$H spectrum of B7. The NMR spectrum was recorded at ~ 500 µM concentration, 50 mM $KH_2PO_4$, pH = 7.5, 50 mM KCl, 1 mM $MgCl_2$, 10% $D_2O$ and 2% DMSO on a 600 MHz Bruker spectrometer equipped with a cryoprobe. The red asterisk indicates the DMSO peak.

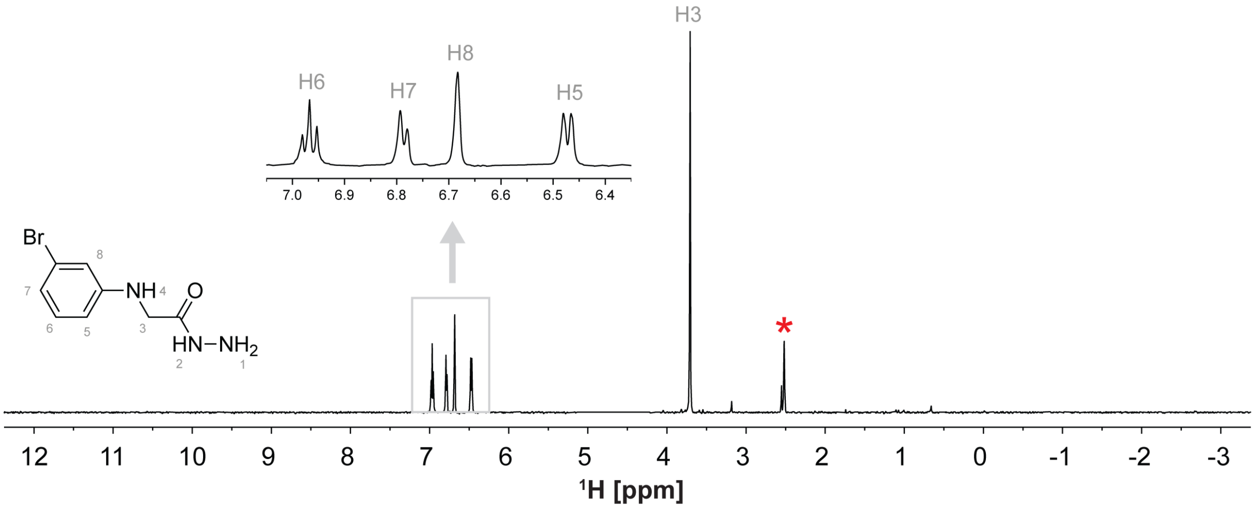


**Figure S18.** Chemical structure and $^1$H spectrum of B8. The NMR spectrum was recorded at ~ 500 µM concentration, 50 mM $KH_2PO_4$, pH = 7.5, 50 mM KCl, 1 mM $MgCl_2$, 10% $D_2O$ and 2% DMSO on a 600 MHz Bruker spectrometer equipped with a cryoprobe. The red asterisk indicates the DMSO peak.

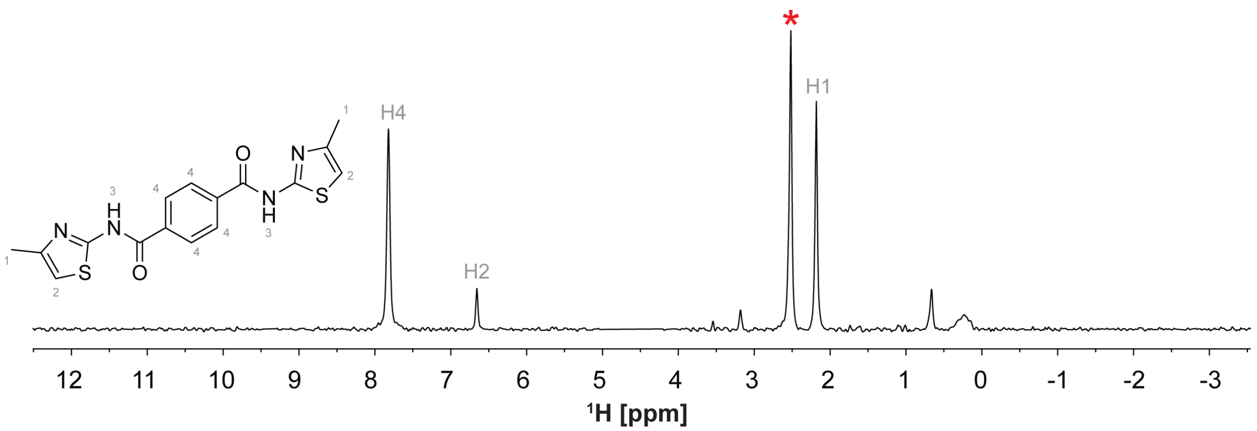


**Figure S19.** Chemical structure and $^1$H spectrum of B9. The NMR spectrum was recorded at ~ 500 µM concentration, 50 mM $KH_2PO_4$, pH = 7.5, 50 mM KCl, 1 mM $MgCl_2$, 10% $D_2O$ and 2% DMSO on a 600 MHz Bruker spectrometer equipped with a cryoprobe. The red asterisk indicates the DMSO peak.

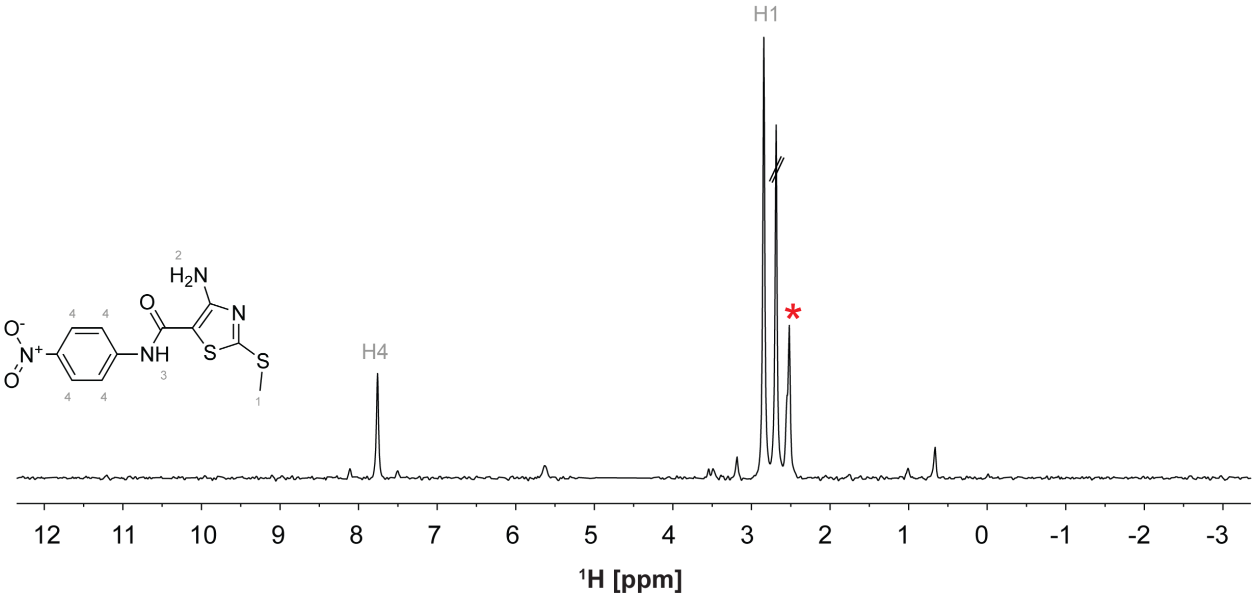


**Figure S20.** Chemical structure and ¹H spectrum of B10. The NMR spectrum was recorded at ~ 500 µM concentration, 50 mM $KH_2PO_4$, pH = 7.5, 50 mM KCl, 1 mM $MgCl_2$, 10% $D_2O$ and 2% DMSO on a 600 MHz Bruker spectrometer equipped with a cryoprobe. The red asterisk indicates the DMSO peak. The extra peak may arise from trace reagent impurities

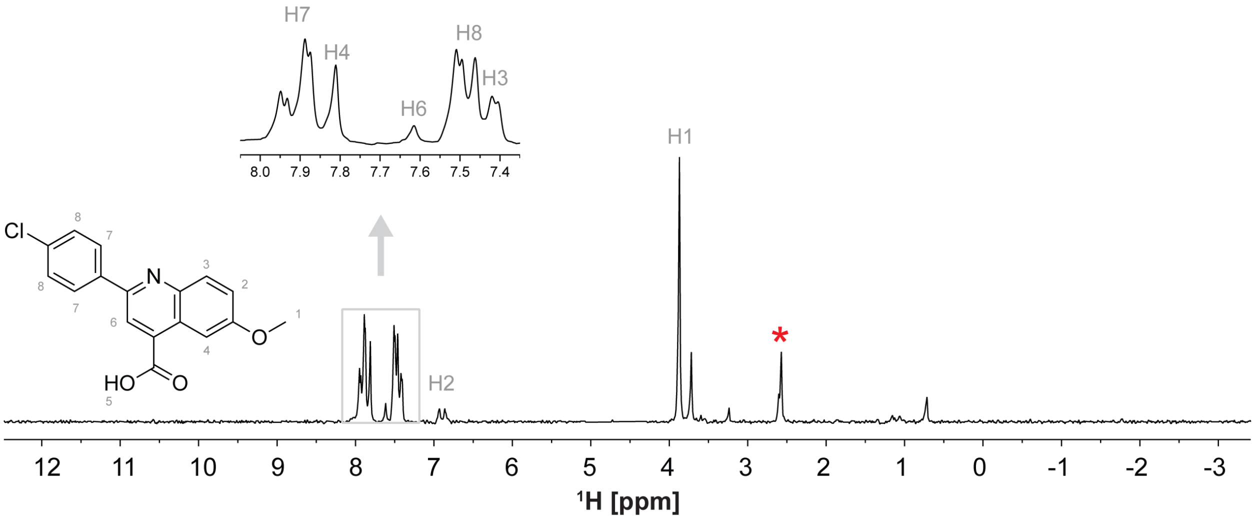


**Figure S21.** Chemical structure and [1]H spectrum of C1. The NMR spectrum was recorded at ~ 500 µM concentration, 50 mM $KH_2PO_4$, pH = 7.5, 50 mM KCl, 1 mM $MgCl_2$, 10% $D_2O$ and 2% DMSO on a 600 MHz Bruker spectrometer equipped with a cryoprobe. The red asterisk indicates the DMSO peak.

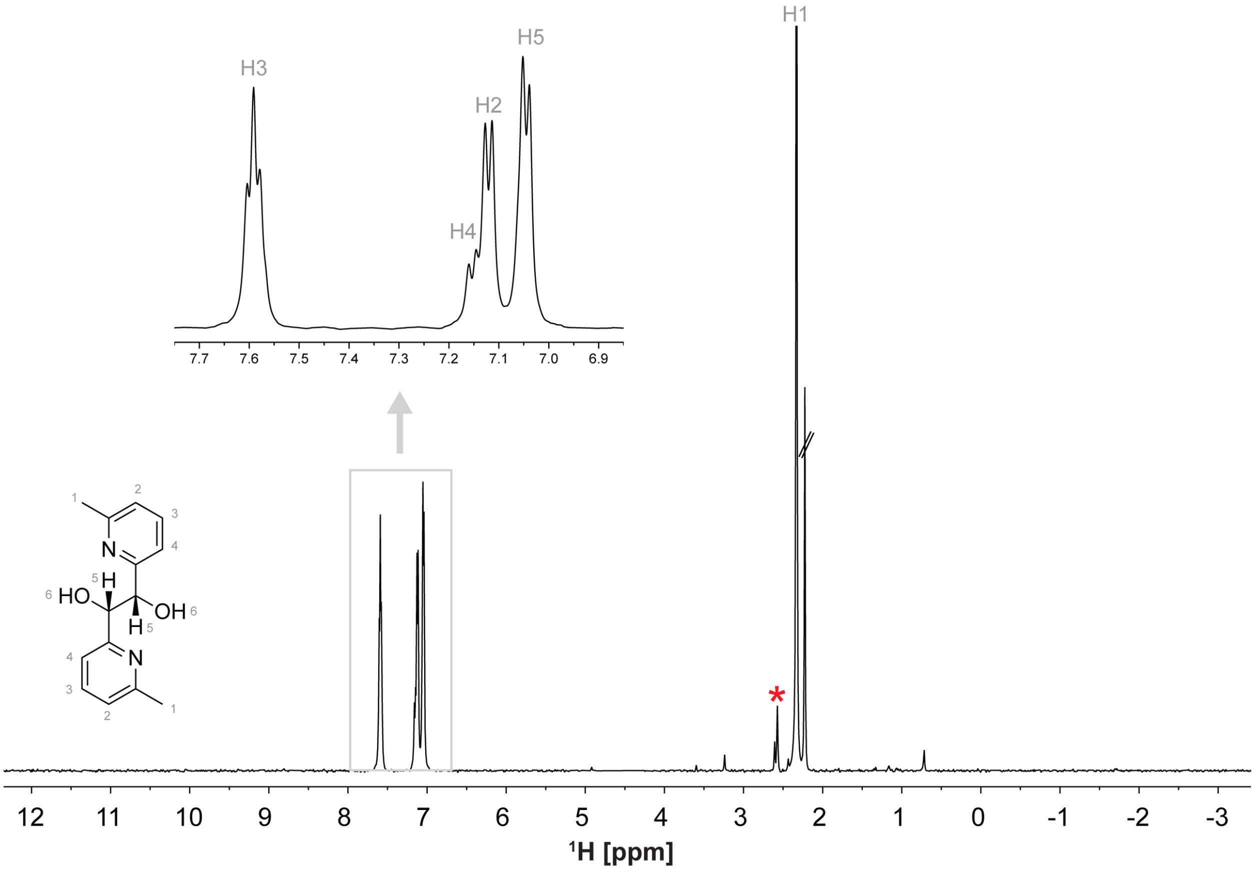


**Figure S22.** Chemical structure and $^{1}$H spectrum of C2. The NMR spectrum was recorded at ~ 500 μM concentration, 50 mM $KH_2PO_4$, pH = 7.5, 50 mM KCl, 1 mM $MgCl_2$, 10% $D_2O$ and 2% DMSO on a 600 MHz Bruker spectrometer equipped with a cryoprobe. The red asterisk indicates the DMSO peak. The extra peak may arise from trace reagent impurities.

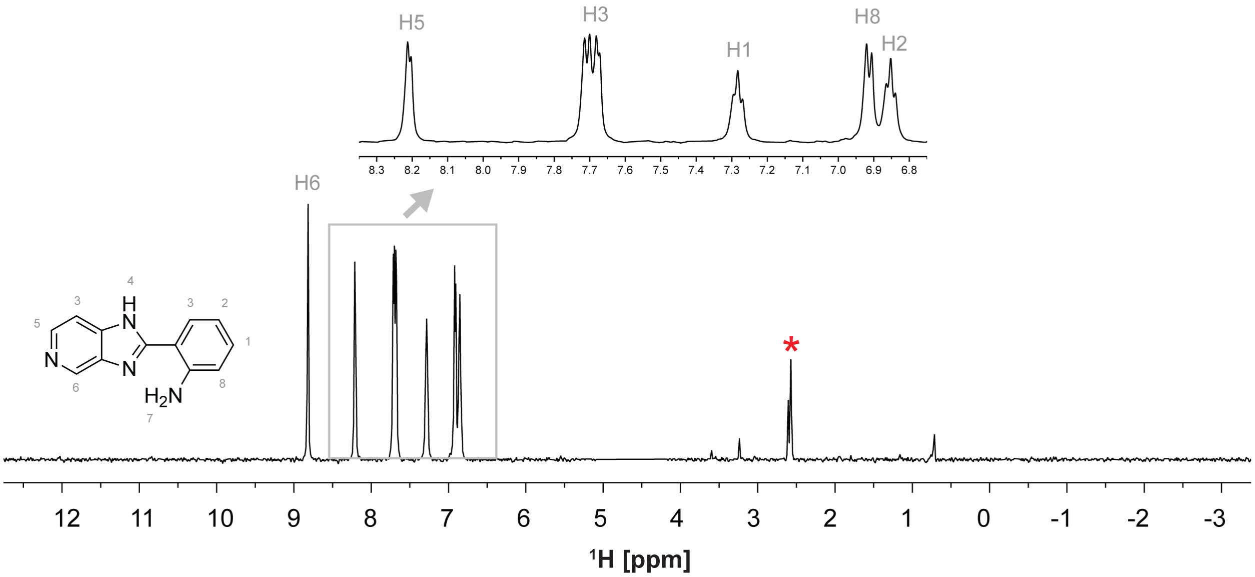


**Figure S23.** Chemical structure and $^1$H spectrum of C2. The NMR spectrum was recorded at ~ 500 µM concentration, 50 mM $KH_2PO_4$, pH = 7.5, 50 mM KCl, 1 mM $MgCl_2$, 10% $D_2O$ and 2% DMSO on a 600 MHz Bruker spectrometer equipped with a cryoprobe. The red asterisk indicates the DMSO peak.

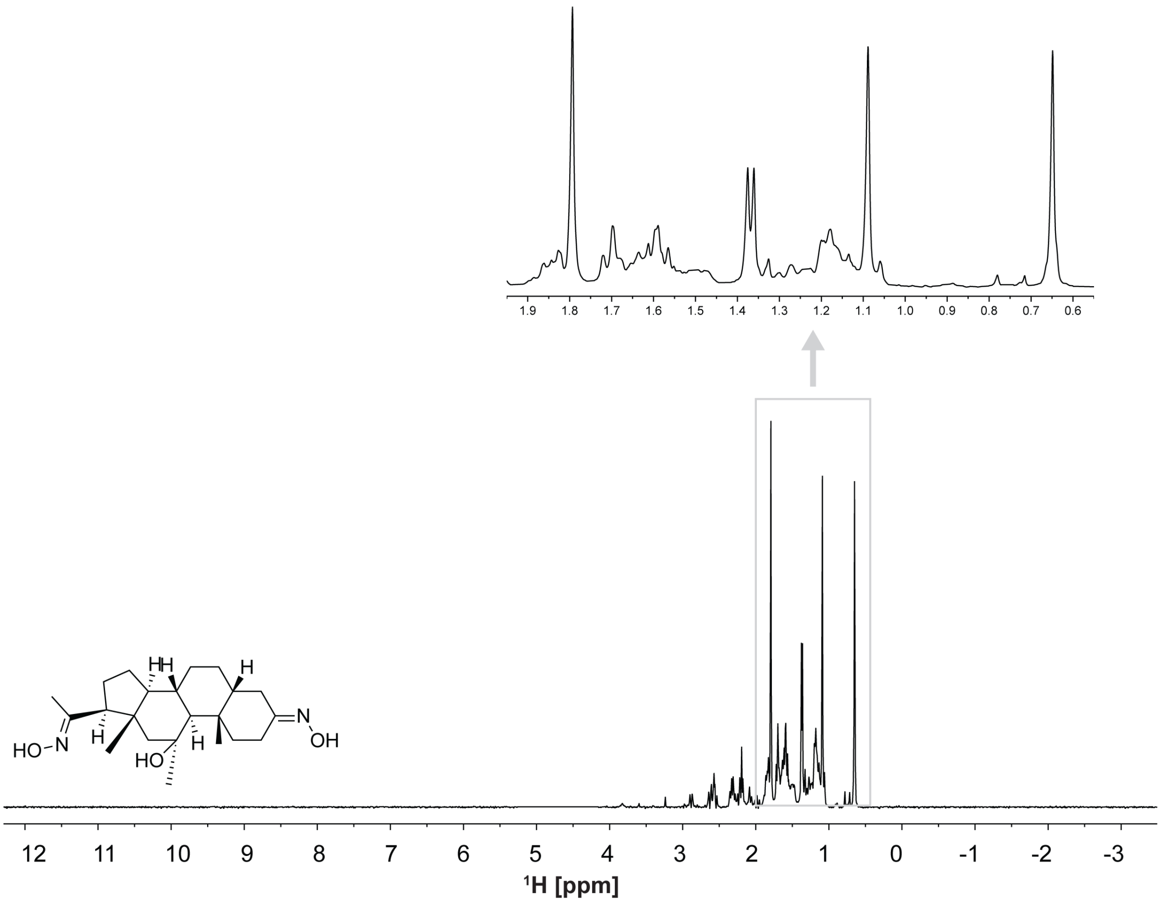


**Figure S24.** Chemical structure and $^1$H spectrum of C6. The NMR spectrum was recorded at ~ 500 µM concentration, 50 mM $KH_2PO_4$, pH = 7.5, 50 mM KCl, 1 mM $MgCl_2$, 10% $D_2O$ and 2% DMSO on a 600 MHz Bruker spectrometer equipped with a cryoprobe. The DMSO peak is overlapped with some signals in C6.

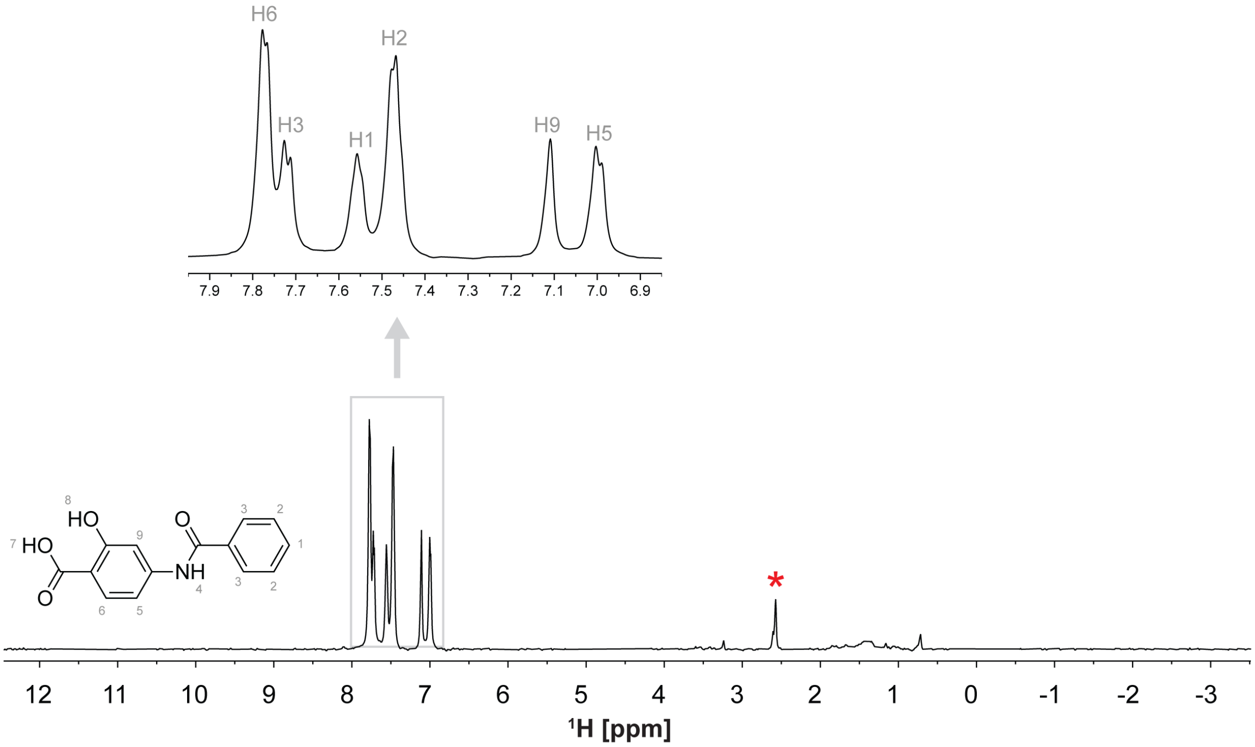


**Figure S25.** Chemical structure and $^{1}$H spectrum of C7. The NMR spectrum was recorded at ~ 500 μM concentration, 50 mM $KH_2PO_4$, pH = 7.5, 50 mM KCl, 1 mM $MgCl_2$, 10% $D_2O$ and 2% DMSO on a 600 MHz Bruker spectrometer equipped with a cryoprobe. The red asterisk indicates the DMSO peak.

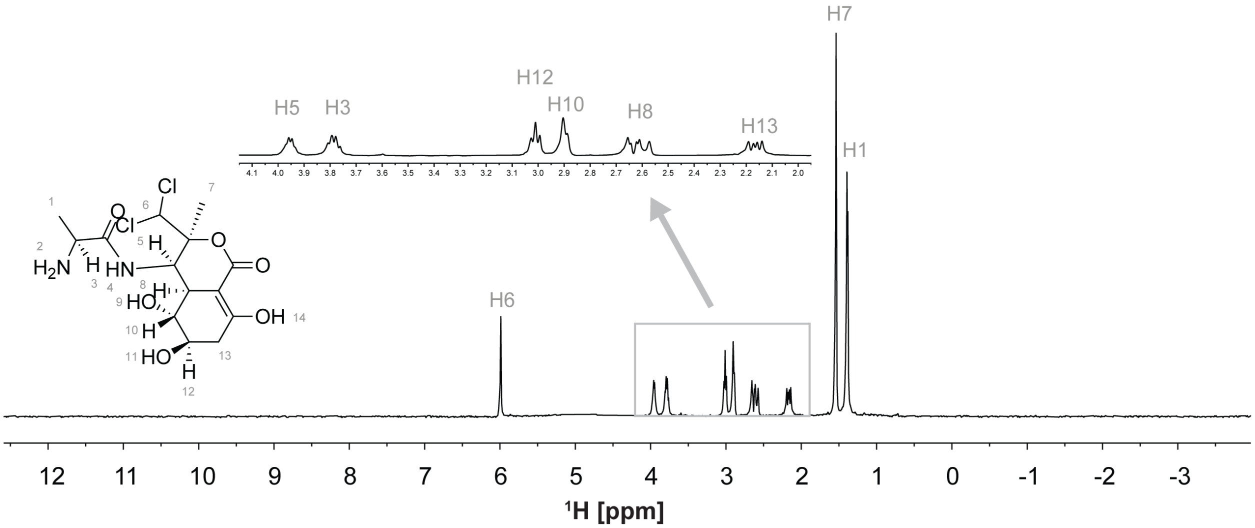


**Figure S26.** Chemical structure and $^1$H spectrum of RA2. The NMR spectrum was recorded at ~ 500 µM concentration, 50 mM $KH_2PO_4$, pH = 7.5, 50 mM KCl, 1 mM $MgCl_2$, 10% $D_2O$ and 2% DMSO on a 600 MHz Bruker spectrometer equipped with a cryoprobe. The DMSO peak is overlapped with some signals in RA2

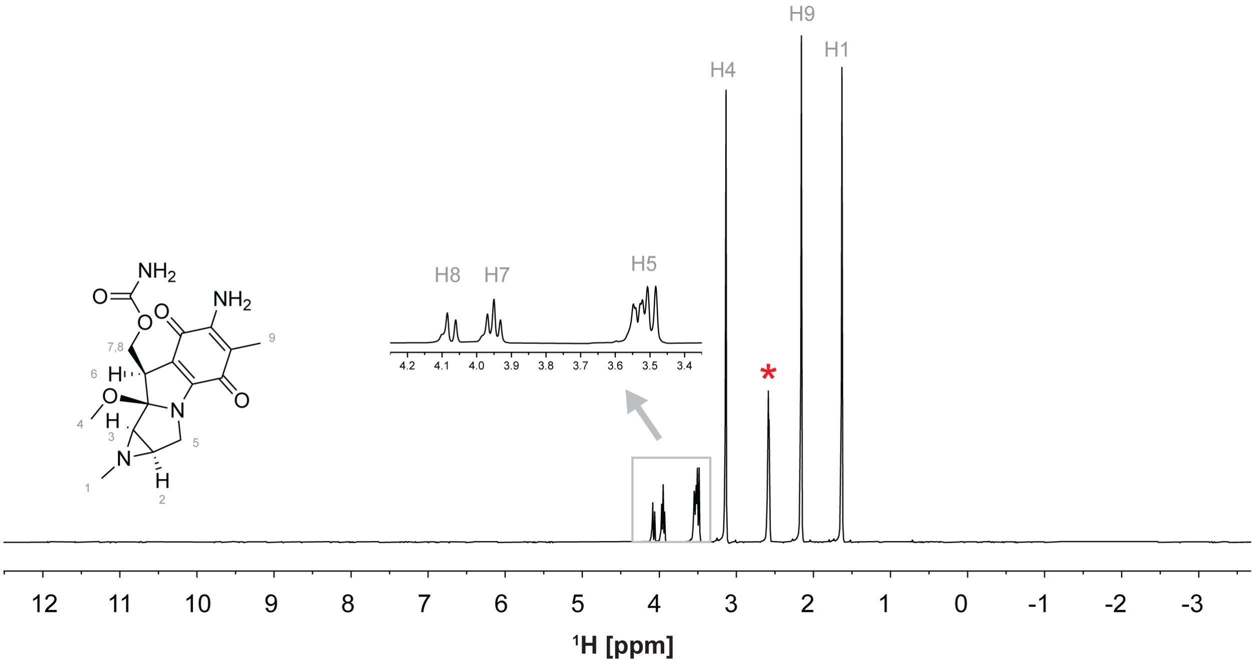


**Figure S27.** Chemical structure and $^1H$ spectrum of RA3. The NMR spectrum was recorded at ~ 500 µM concentration, 50 mM $KH_2PO_4$, pH = 7.5, 50 mM KCl, 1 mM $MgCl_2$, 10% $D_2O$ and 2% DMSO on a 600 MHz Bruker spectrometer equipped with a cryoprobe. The red asterisk indicates the DMSO peak.

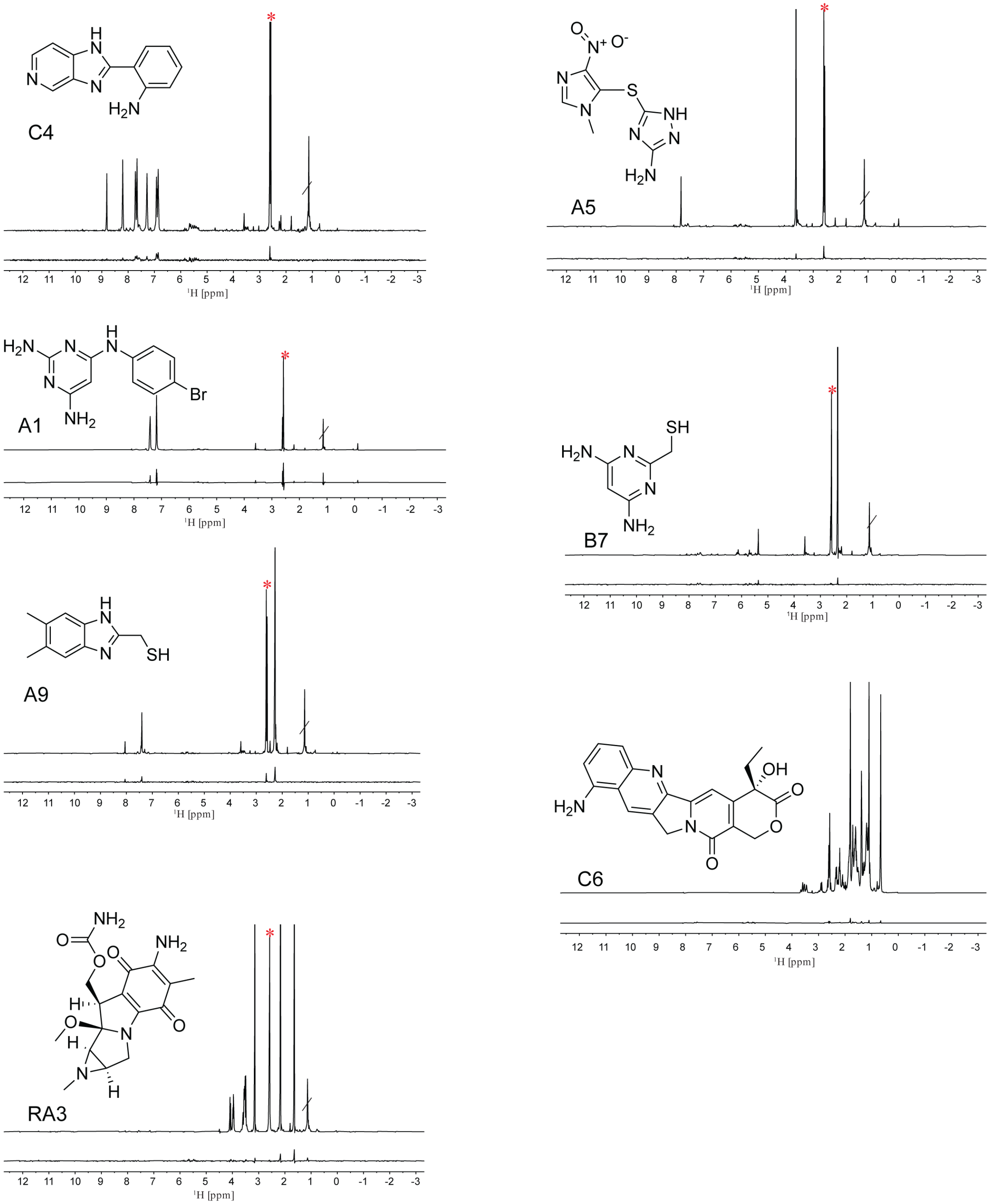


**Figure S28.** STD-NMR identified ‘hits’ of the miR-31 hairpin. The ‘off’ resonance spectrum (top) and difference spectrum (bottom) as well as chemical structures of the small molecules are shown.

The red asterisk present in all spectra except for C6 indicates the DMSO peak.

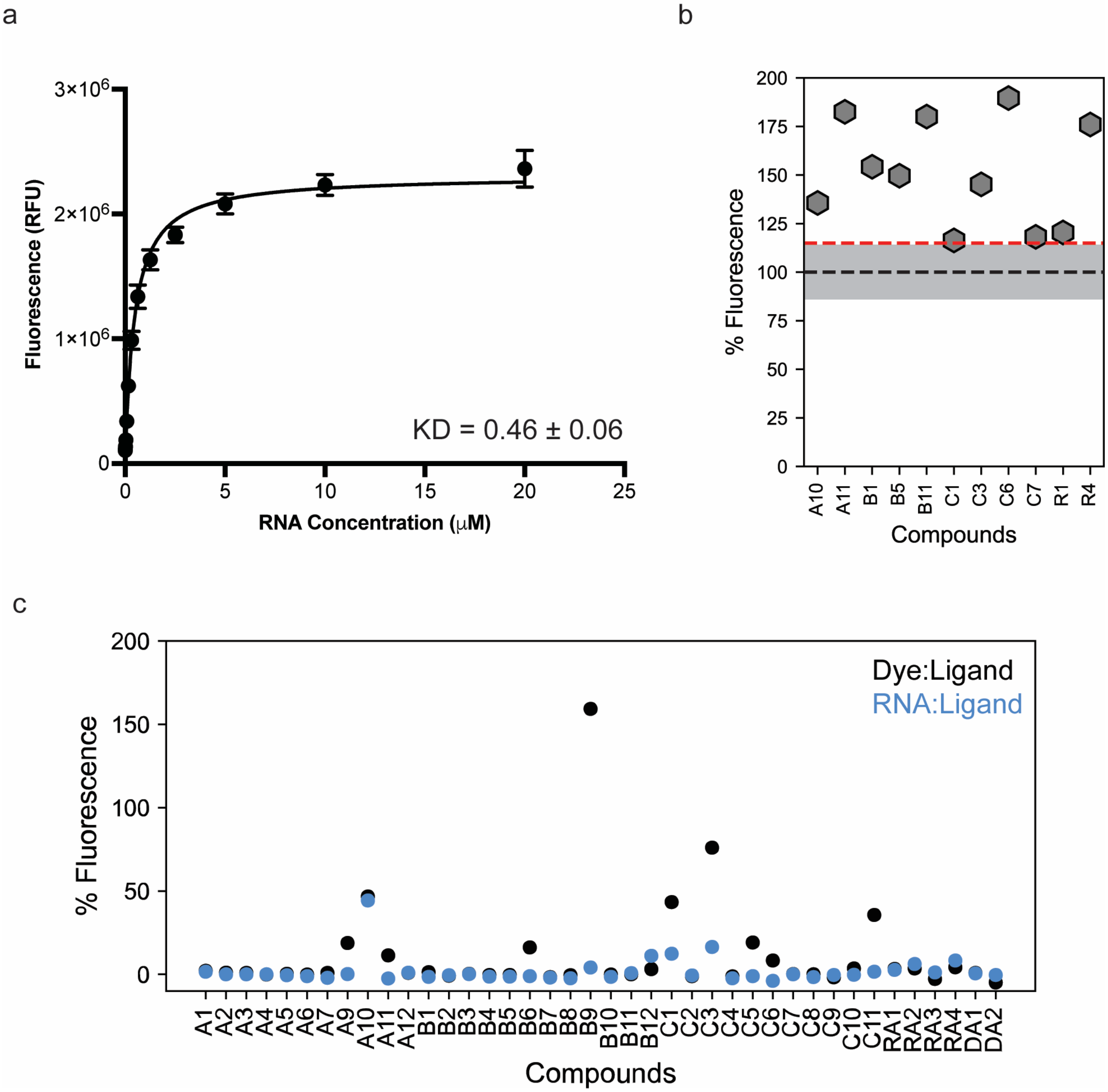


**Figure S29.** Flourescence indicator displacement assay. (a) Affinity of TO-PRO-1 to the miR-31 hairpin. (b) Percentage fluorescence change in RNA-dye complex by small molecules that resulted in significant fluorescence increase. (c) Percentage fluorescence change in RNA-ligand when compared to RNA-buffer (blue) and Dye-ligand when compared to Dye-buffer alone.

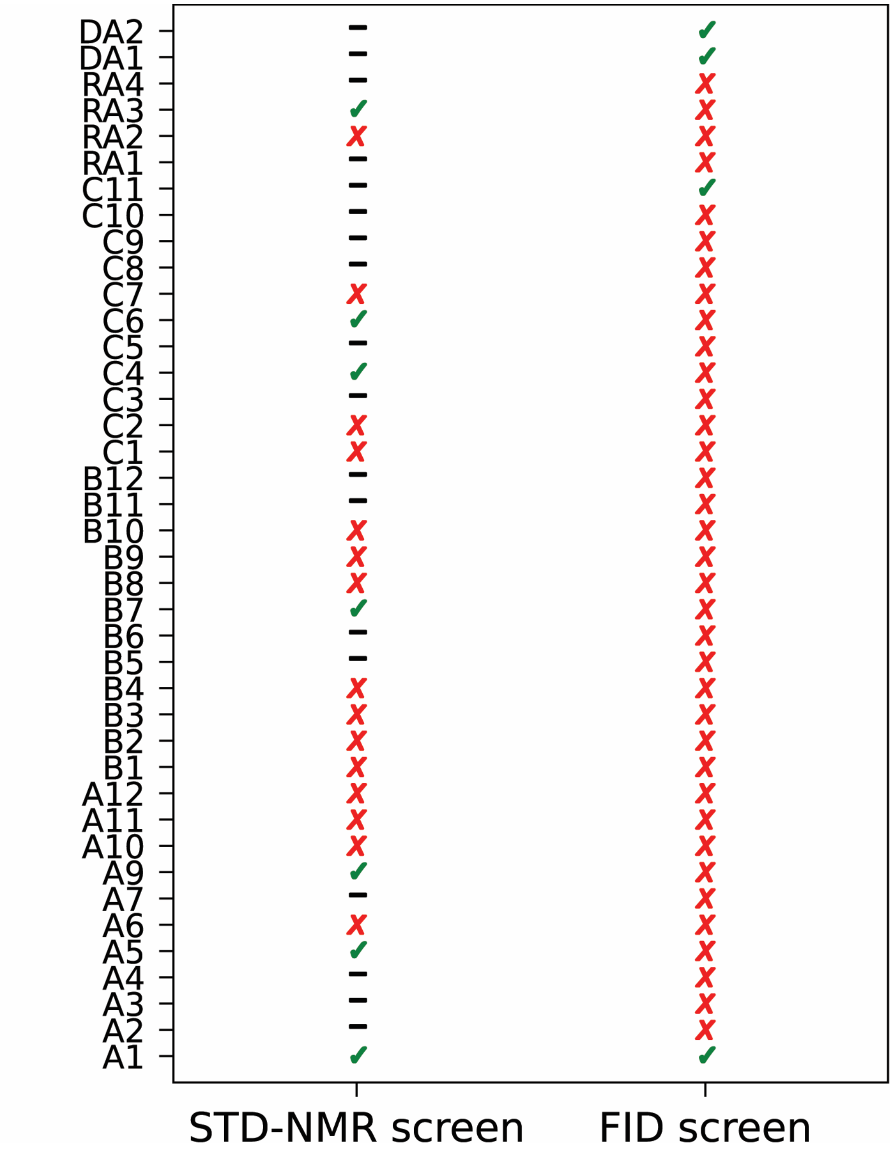


**Figure S30.** Summary of screening results from two independent assays. Lanes indicated with a black dash (-) denote no data, a green check mark denotes a hit and a red X indicates a non-hit.

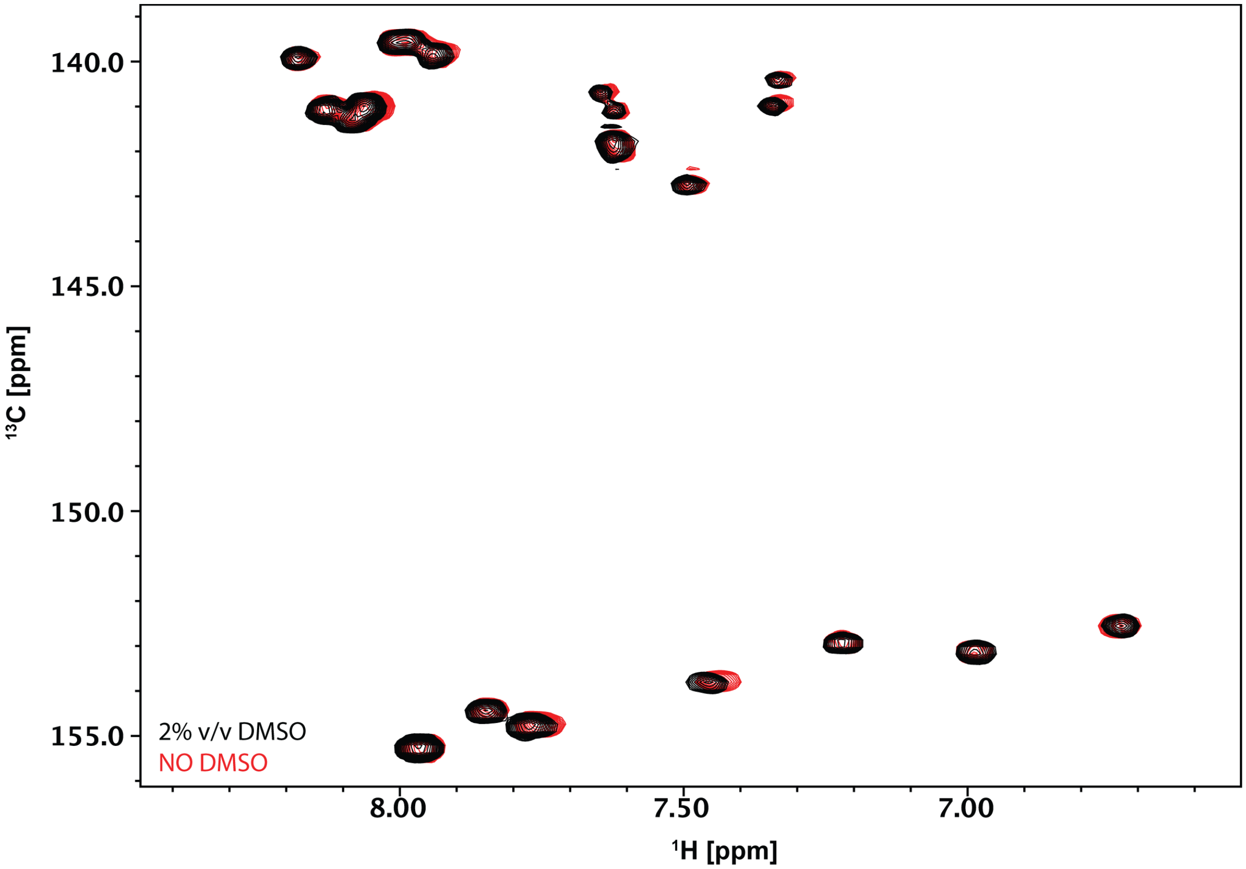


**Figure S31.** Overlayed $^{1}H$-$^{13}C$ HSQC spectra of $^{13}C/^{15}N$ adenosine cytosine labeled miR-31 hairpin at 0% (v/v) DMSO (red) and 2% (v/v) DMSO (black).

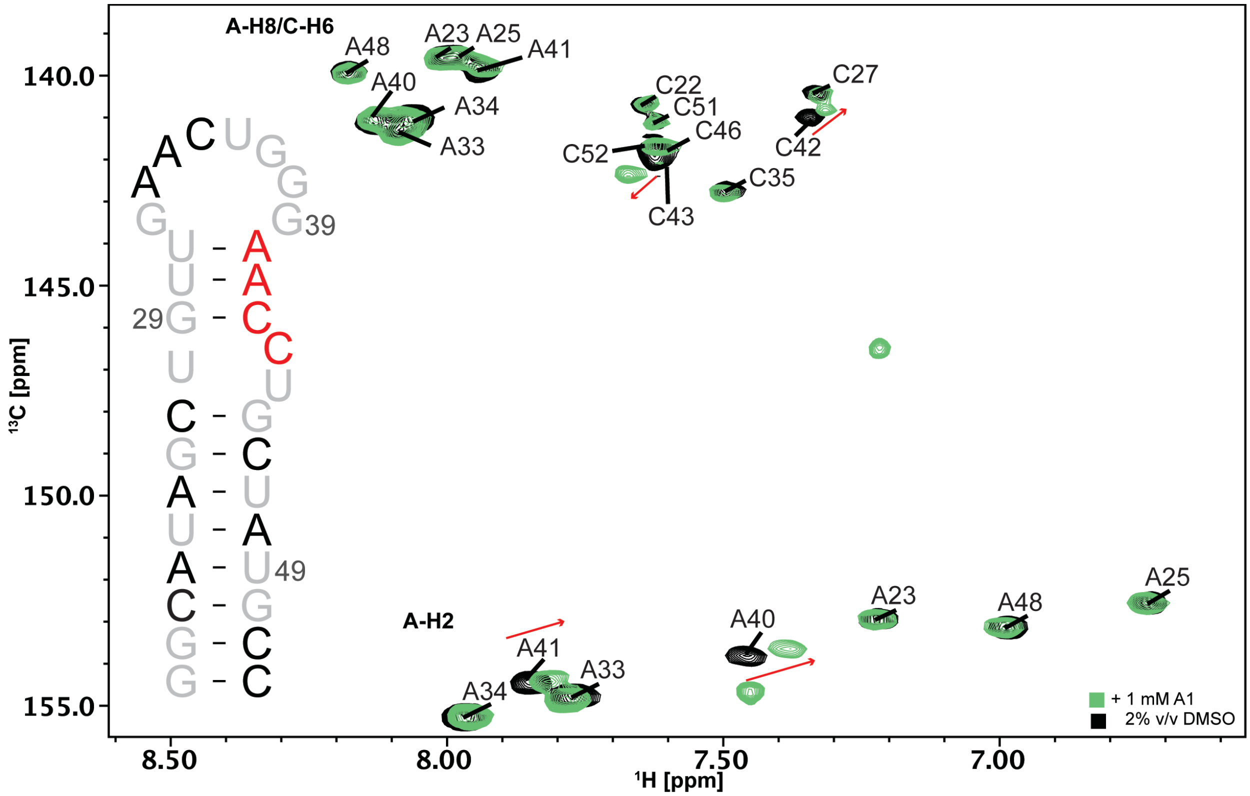


**Figure S32.** Overlayed $^1$H-$^{13}$C HSQC spectra of $^{13}$C/$^{15}$N adenosine cytosine labeled miR-31 hairpin in 2% (v/v) DMSO (black) and 1 mM A1 (green). Significant perturbations are mapped on the secondary structure of the miR-31 hairpin.

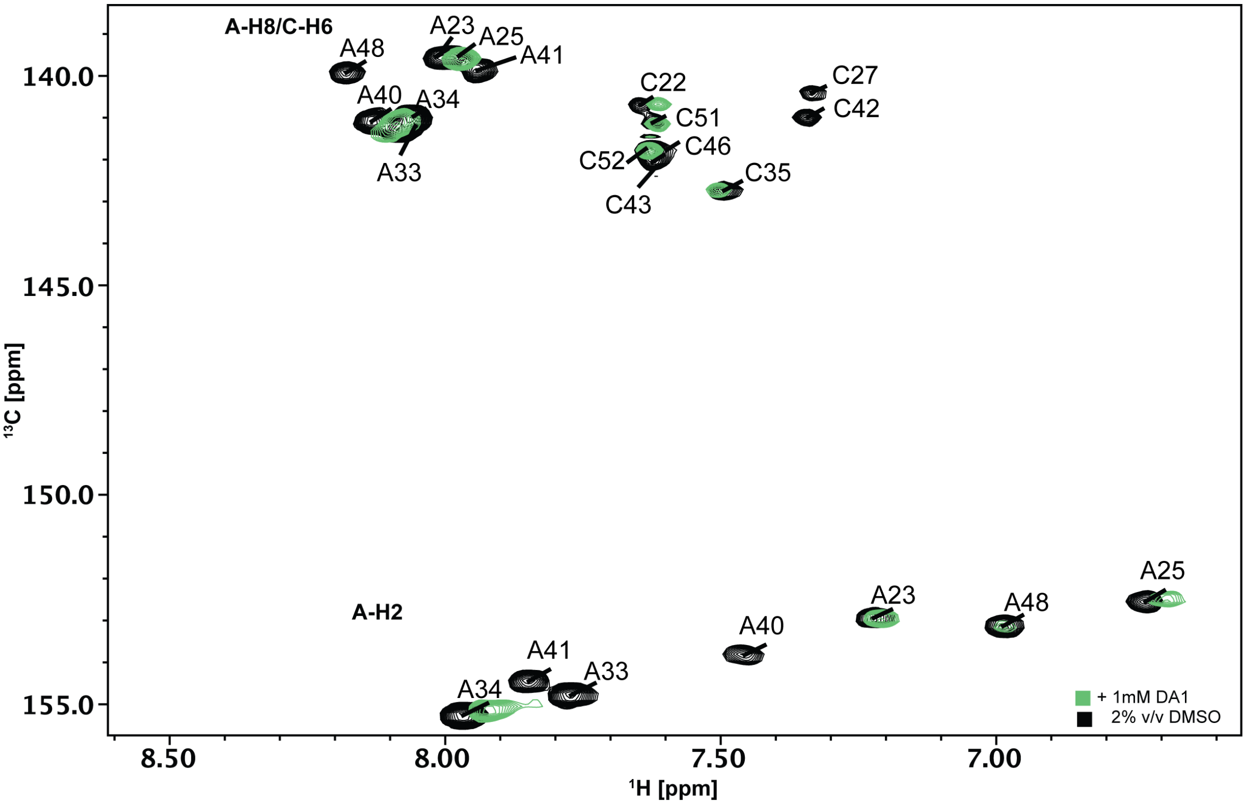


**Figure S33.** Overlayed $^{1}H$-$^{13}C$ HSQC spectra of $^{13}C/^{15}N$ adenosine cytosine labeled miR-31 hairpin in 2% (v/v) DMSO (black) and ~1 mM DA1 (green).

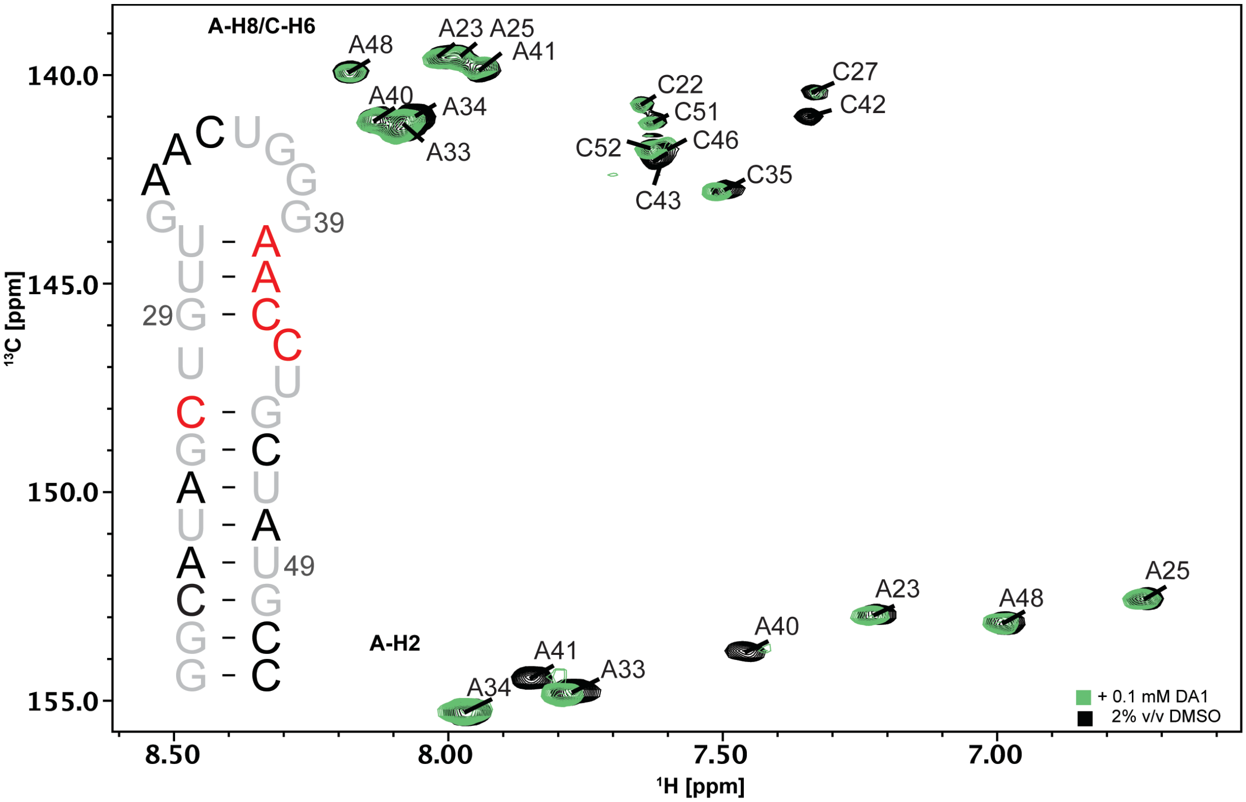


**Figure S34.** Overlayed $^{1}H$-$^{13}C$ HSQC spectra of $^{13}C/^{15}N$ adenosine cytosine labeled miR-31 hairpin in 2% (v/v) DMSO (black) and 0.1 mM DA1 (green). Significant perturbations are mapped on the secondary structure of the miR-31 hairpin.

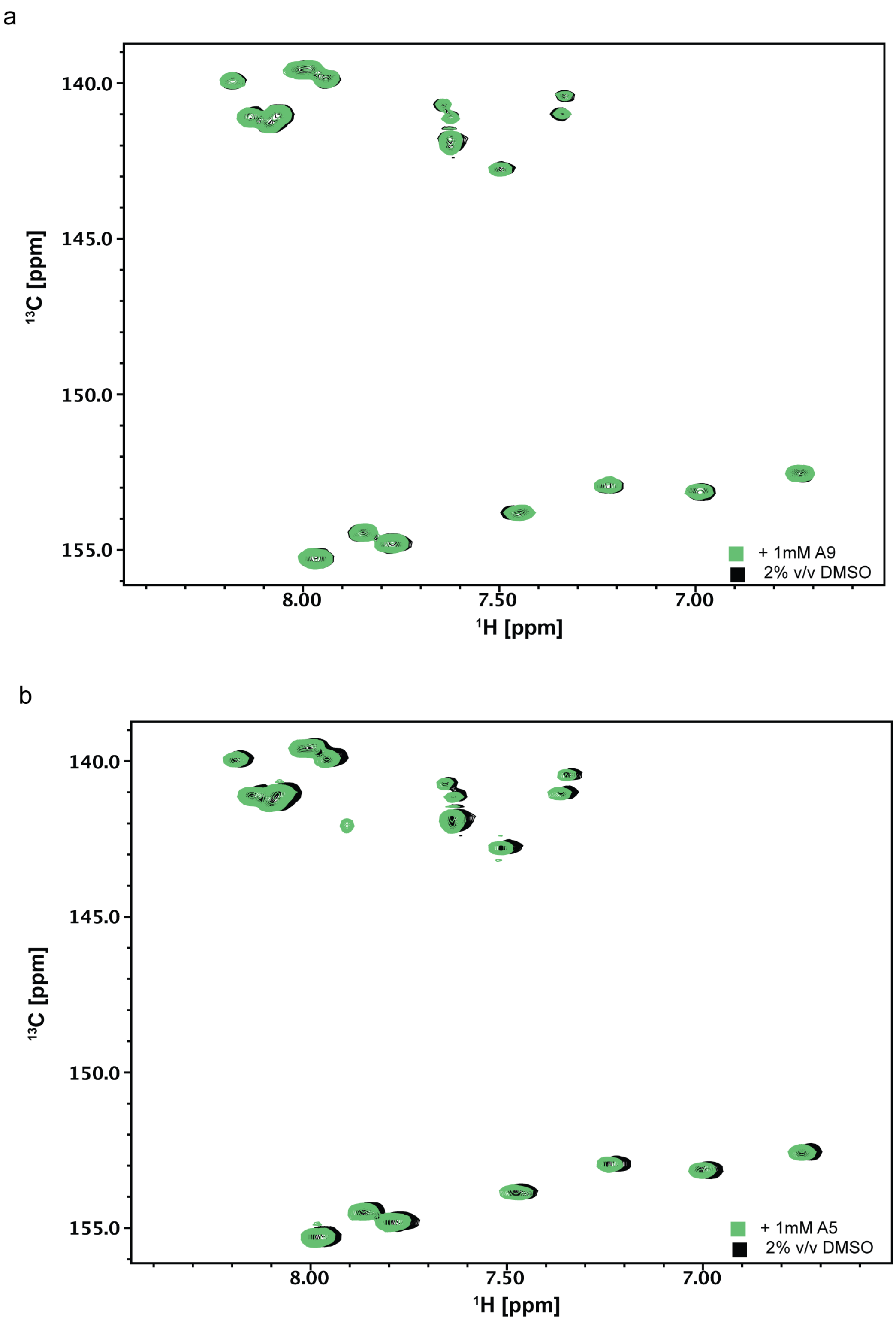


**Figure S35.** (a-b) Overlayed $^{1}$H-$^{13}$C HSQC spectra of $^{13}$C/$^{15}$N adenosine cytosine labeled miR-31 hairpin in 2% (v/v) DMSO (black) and (a) 1 mM A9 (green) or (b) 1 mM A5 (green).

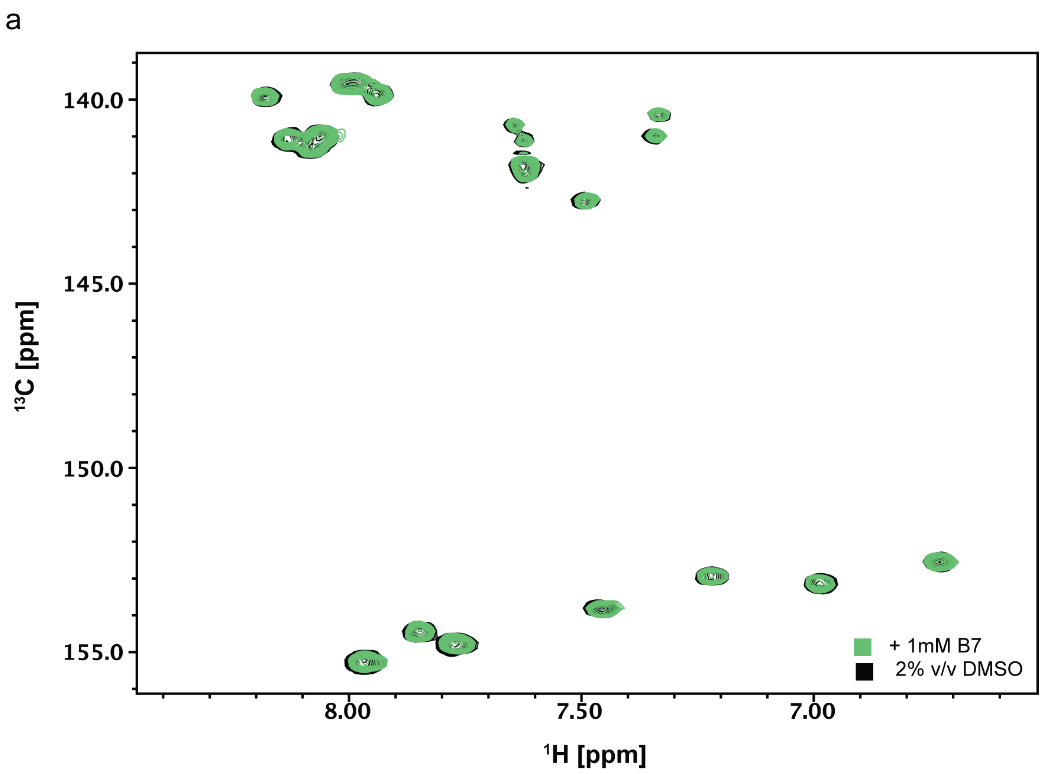


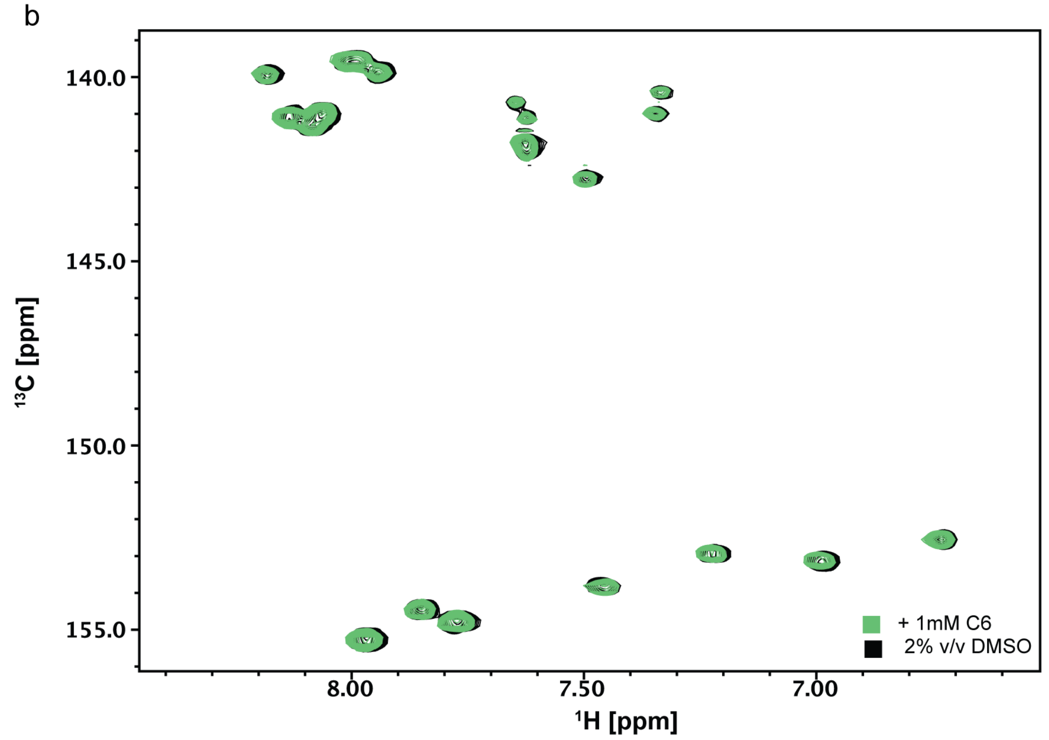


**Figure S36.** (a-b) Overlayed $^{1}H$-$^{13}C$ HSQC spectra of $^{13}C/^{15}N$ adenosine cytosine labeled miR-31 hairpin in 2% (v/v) DMSO (black) and (a) 1 mM B7 (green) or (b) 1 mM C6 (green).

a

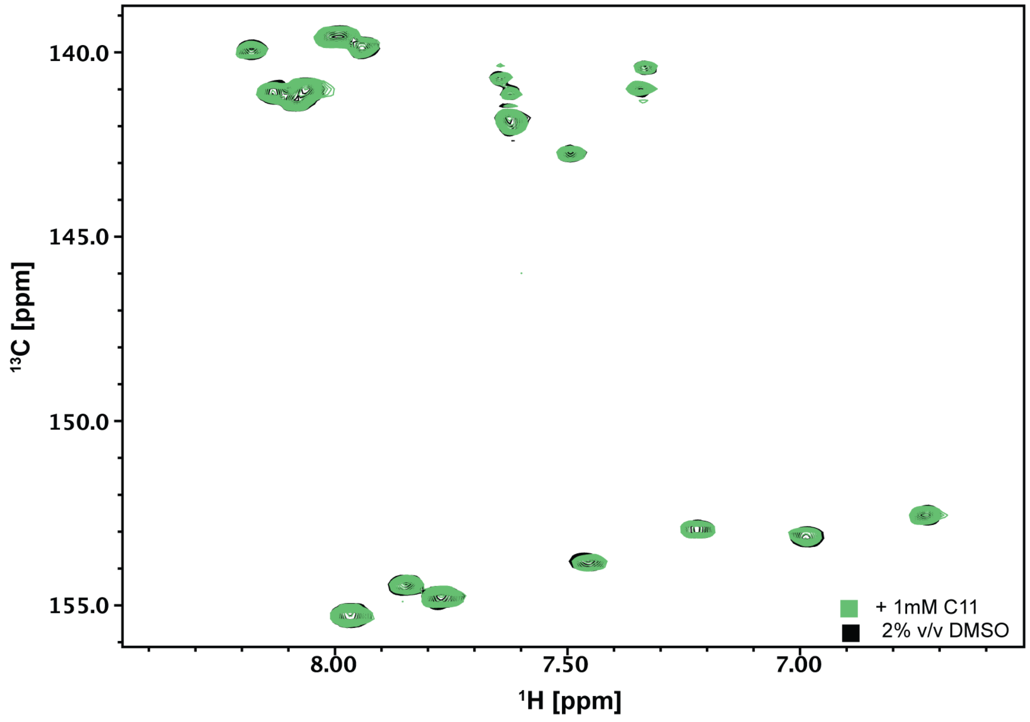


b

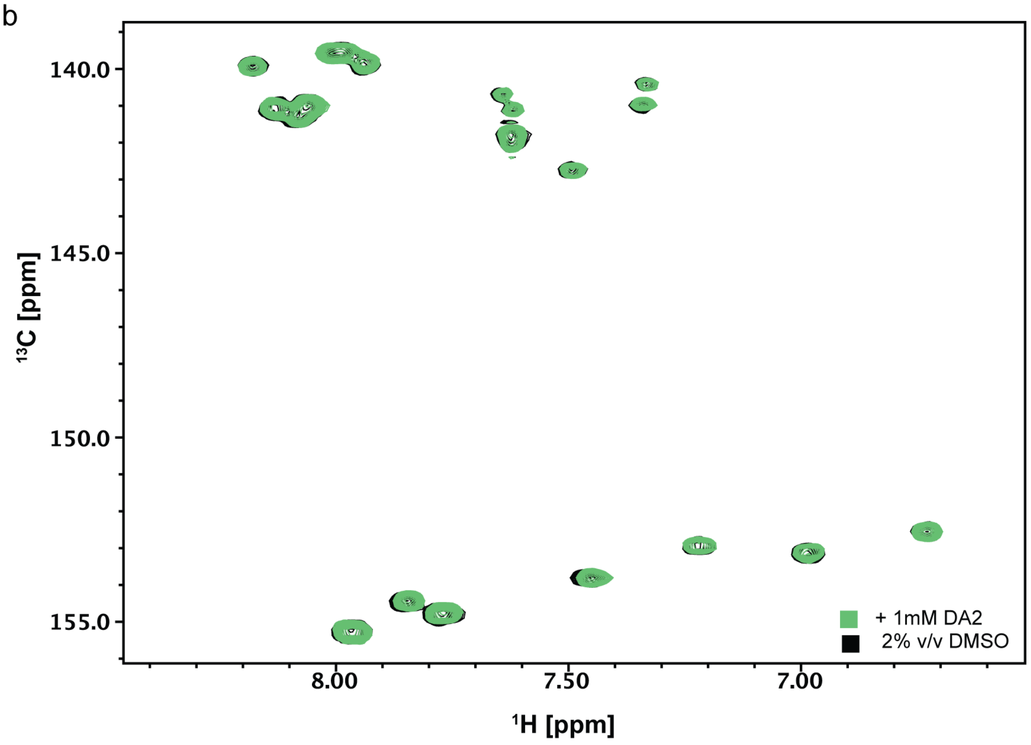


**Figure S37.** (a-b) Overlayed $^{1}$H-$^{13}$C HSQC spectra of $^{13}$C/$^{15}$N adenosine cytosine labeled miR-31 hairpin in 2% (v/v) DMSO (black) and (a) ~1 mM C11 (green) or (b) ~1 mM DA2 (green).

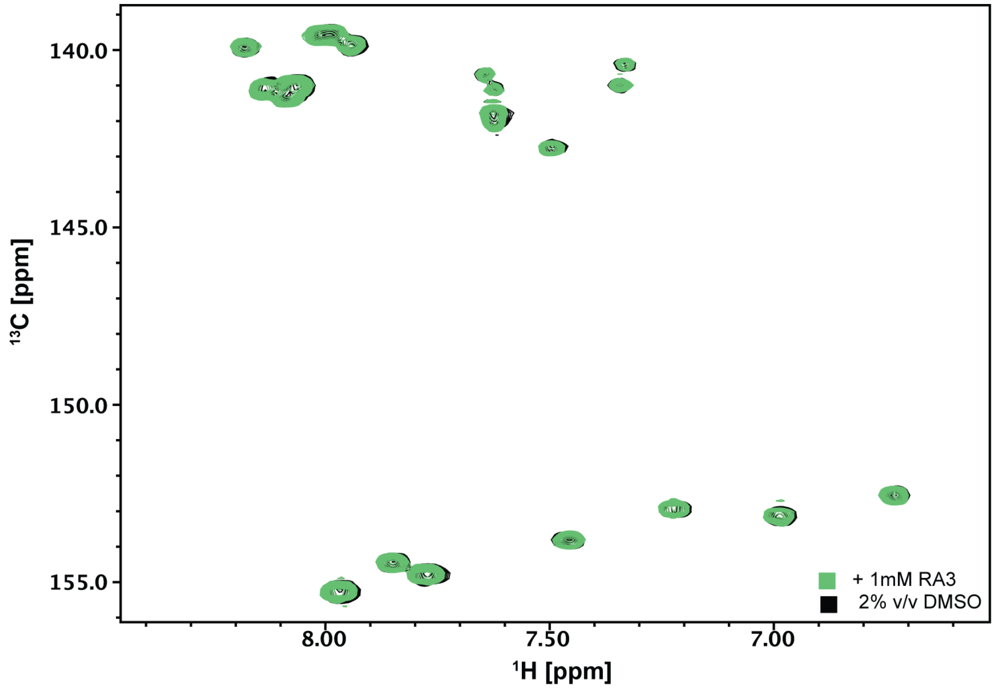


**Figure S38.** Overlayed $^{1}H$-$^{13}C$ HSQC spectra of $^{13}C$/$^{15}N$ adenosine cytosine labeled miR-31 hairpin in 2% (v/v) DMSO (black) and ~1 mM RA3 (green).

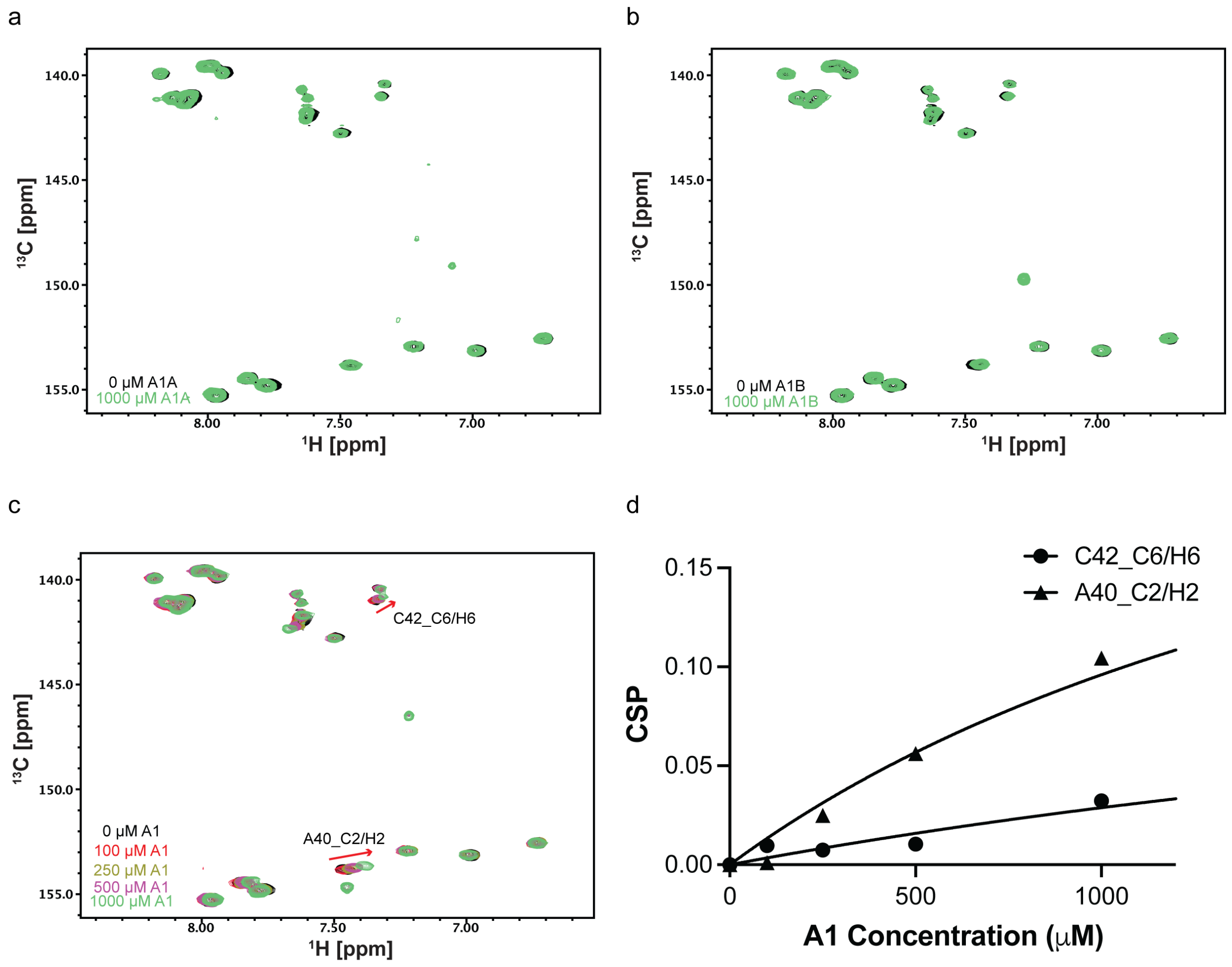


**Figure S39.** Binding affinity estimation of A1 and miR-31 hairpin. (a-b) Overlayed $^{1}$H-$^{13}$C spectra of miR-31 hairpin with (a) no ligand (black) and 1 mM A1A green), (b) no ligand (black) and 1 mM A1B (green). (c) Overlayed $^{1}$H-$^{13}$C spectra of miR-31 hairpin at different A1 concentrations. (d) Fits of the chemical shift perturbations as function of A1 concentration.

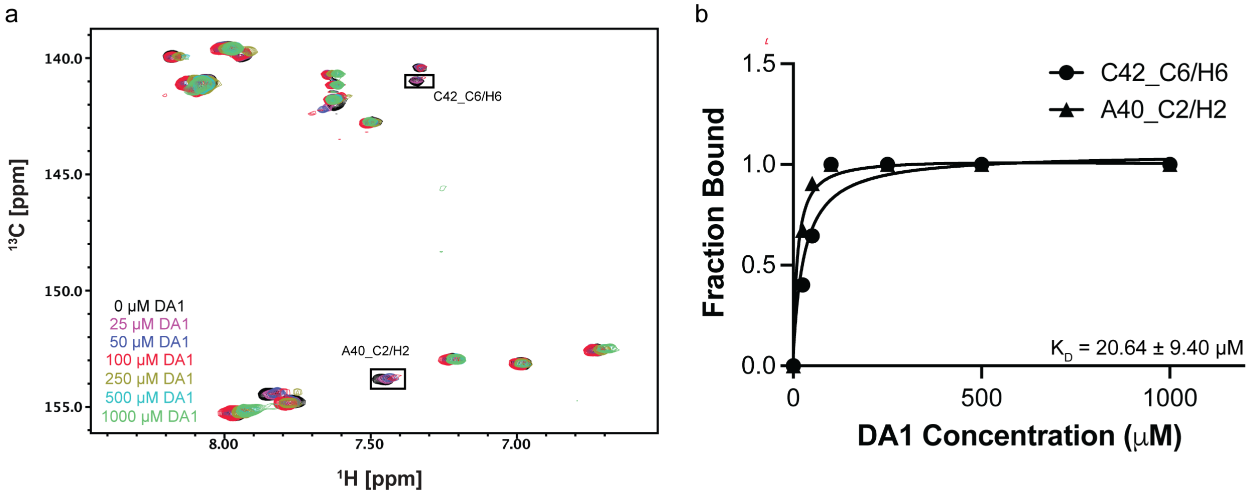


**Figure S40.** Binding affinity estimation of DA1 and miR-31 hairpin. (a) Overlayed $^{1}H$-$^{13}C$ spectra of miR-31 hairpin at different DA1 concentrations. (b) Fits of the chemical shift perturbations as function of DA1 concentration.
